# Dual Role of Cell-Cell Adhesion In Tumor Suppression and Proliferation Due to Collective Mechanosensing

**DOI:** 10.1101/683250

**Authors:** Abdul N Malmi-Kakkada, Xin Li, Sumit Sinha, D. Thirumalai

## Abstract

It is known that mechanical interactions couple a cell to its neighbors, enabling a feedback loop to regulate tissue growth. However, the interplay between cell-cell adhesion strength, local cell density and force fluctuations in regulating cell proliferation is poorly understood. Here, we show that spatial variations in the tumor growth rates, which depend on the location of cells within tissue spheroids, are strongly influenced by cell-cell adhesion. As the strength of the cell-cell adhesion increases, intercellular pressure initially decreases, enabling dormant cells to more readily enter into a proliferative state. We identify an optimal cell-cell adhesion regime where pressure on a cell is a minimum, allowing for maximum proliferation. We use a theoretical model to validate this novel collective feedback mechanism coupling adhesion strength, local stress fluctuations and proliferation. Our results predict the existence of a non-monotonic proliferation behavior as a function of adhesion strength, consistent with experimental results. Several experimental implications of the proposed role of cell-cell adhesion in proliferation are quantified, making our model predictions amenable to further experimental scrutiny. We show that the mechanism of contact inhibition of proliferation, based on a pressure-adhesion feedback loop, serves as a unifying mechanism to understand the role of cell-cell adhesion in proliferation.

## I. INTRODUCTION

Adhesive forces between cells, mediated by cadherins, play a critical role in morphogenesis, tissue healing, and tumor growth [1, 2]. In these processes, the collective cell properties are influenced by how cells adhere to one another, enabling cells to communicate through mechanical forces [3, 4]. Amongst the family of cadherins, E-cadherin is the most abundant, found in most metazoan cohesive tissues [5]. E-cadherin transmembrane proteins facilitate intercellular bonds through the interaction of extracellular domains on neighboring cells. The function of cadherins was originally appreciated through their role in cell aggregation during morphogenesis [6, 7]. Mechanical coupling between the cortical cytoskeleton and cell membrane is understood to involve the cadherin cytoplasmic domain [8]. Forces exerted across cell-cell contacts is transduced between cadherin extracellular domain and the cellular cytoskeletal machinery through the cadherin/catenin complex [9]. Therefore, to understand how mechanical forces control the spatial organization of cells within tissues and impact proliferation, the role of adhesion strength at cell-cell contacts need to be elucidated.

Together with cell-cell adhesion, cell proliferation control is of fundamental importance in animal and plant development, regeneration, and cancer progression [10, 11]. Spatial constraints due to cell packing or crowding are known to affect cell proliferation [12–17]. The spatiotemporal arrangement of cells in response to local stress field fluctuations, arising from intercellular interactions, and how it feeds back onto cell proliferation remains unclear. Indeed, evidence so far based on experimental and theoretical studies on the mechanism underlying the crosstalk between the strength of cell-cell adhesion and proliferation, invasion and drug resistance is not well understood [15, 18–21].

We briefly discuss two seemingly paradoxical roles of E-cadherin in proliferation. E-cadherin depletion lead cells to adopt a mesenchymal morphology characterized by enhanced cell migration and invasion [22, 23] while increased expression of E-cadherin in cell lines with minimal expression reverses highly proliferative and invasive phenotypes [24, 25]. Besides suppressing tumor growth, there is also evidence that cadherin expression can lead to tumor proliferation. We detail the dual role that E-cadherin plays below.

### E-cadherin downregulation and tumor progression through the EMT mechanism

Loss of E-cadherin expression is related to the epithelial-to-mesenchymal transition (EMT), observed during embryogenesis [6, 26], tumor progression and metastasis [27, 28]. EMT results in the transformation of epithelial cells into a mesenchymal phenotype, where adhesive strength between the cells is significantly decreased, and drives tumor invasiveness and cell migration in epithelial tumors. In this ‘canonical’ picture, down regulation of cell-cell adhesion contributes to cancer progression [29].

### E-cadherin upregulation may promote tumor progression

In contrast to the reports cited above, others argue that EMT may not be required for cancer metastasis [30–32]. In fact, E-cadherin may facilitate collective cell migration that potentiates invasion and metastasis [33, 34]. Increased E-cadherin expression is deemed necessary for the progression of aggressive tumors such as inflammatory breast cancer (IBC) [35] and a glioblastoma multiforme (GBM) subtype [36, 37], and invasive ductal carcinomas [38]. In multiple GBM cell lines, increased E-cadherin expression positively correlated with tumor growth and invasiveness [36]. In normal rat kidney-52E (NRK-52E) and non-tumorigenic human mammary epithelial cells (MCF-10A), E-cadherin engagement stimulated a peak in proliferation capacity through Rac1 activation [39]. By culturing cells in micro-fabricated wells to control for cell spreading on 2D substrates, VE-cadherin mediated contact with neighboring cells showed enhanced proliferation [40]. The dual role of cell-cell adhesion, suppressing proliferation in some cases and promoting it in other instances, therefore warrants further investigation. The contrasting scenario raise unanswered questions that are amenable to analyses using relatively simple models of tumor growth: (1) How does the magnitude of forces exerted by cells on one another influence their overall growth and proliferation? (2) Can a minimal physical model capture the role of cell-cell adhesion in suppressing and enhancing tumor growth? Here we use simulations and theoretical arguments to establish a non-monotonic dependence between cell-cell adhesion strength (*f*^*ad*^; depth of attractive interaction between cells) and proliferation capacity. While the parameter *f*^*ad*^ is a proxy for cell-cell adhesion strength representing E-cadherin expression, we note that adhesion strength may also increase due to cadherin clustering [41], increasing time of contact between cells [41], or through “mechanical polarization”, where cells reorganize their adhesive and cytoskeletal machinery to suppress actin density along cell-cell contact interfaces [42, 43]. We show that cell proliferation increases as *f*^*ad*^ increases, and reaches a maximum at a critical value, 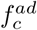. In other words, increasing cell-cell adhesion from low levels causes the tumor proliferative capacity to increase. We identify an intermediate level of cell-cell adhesion where invasiveness and proliferation are maximized. As *f*^*ad*^ is increased beyond 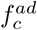, proliferation capacity is suppressed. The non-monotonic dependence of proliferation on *f*^*ad*^ qualitatively explains the dual role of cell-cell adhesion, as we explain below. By building on the integral feedback mechanism coupling cell dormancy and local pressure [15], we suggest a physical pressure based formalism for the effect of cell-cell adhesion on cell proliferation. In particular, we elucidate the role of cell-cell contact, nearest neighbor packing and the onset of force dependent cell growth inhibition in influencing cell proliferation.

## RESULTS

### Cell-Cell Adhesion Strength and Feedback On Cell Proliferation

Cell dynamics within proliferating tissues is a complex process, where the following contributions at the minimum are coupled: (i) cell-cell repulsive and adhesive forces, (ii) cell dynamics due to growth, and (iii) cell division and apoptosis. Stochastic cell growth leads to dynamic variations in the cell-cell forces while cell division and apoptosis induce temporal rearrangements in the cell positions and packing. Cell-cell adhesion strength, dictated by *f*^*ad*^ (see Appendix A Eq. A3), leads to experimentally measurable effects on the spatial arrangement of cells (see Fig. 1a), quantified by the angle, *β*, and the length of contact, *l* _*c*_ between the cells. The angle, *β*, should decrease as a function of *f*^*ad*^ while *l*_*c*_ should increase [43] (see Appendix A Figs. 6a and 6b for further details).

**FIG. 1.**
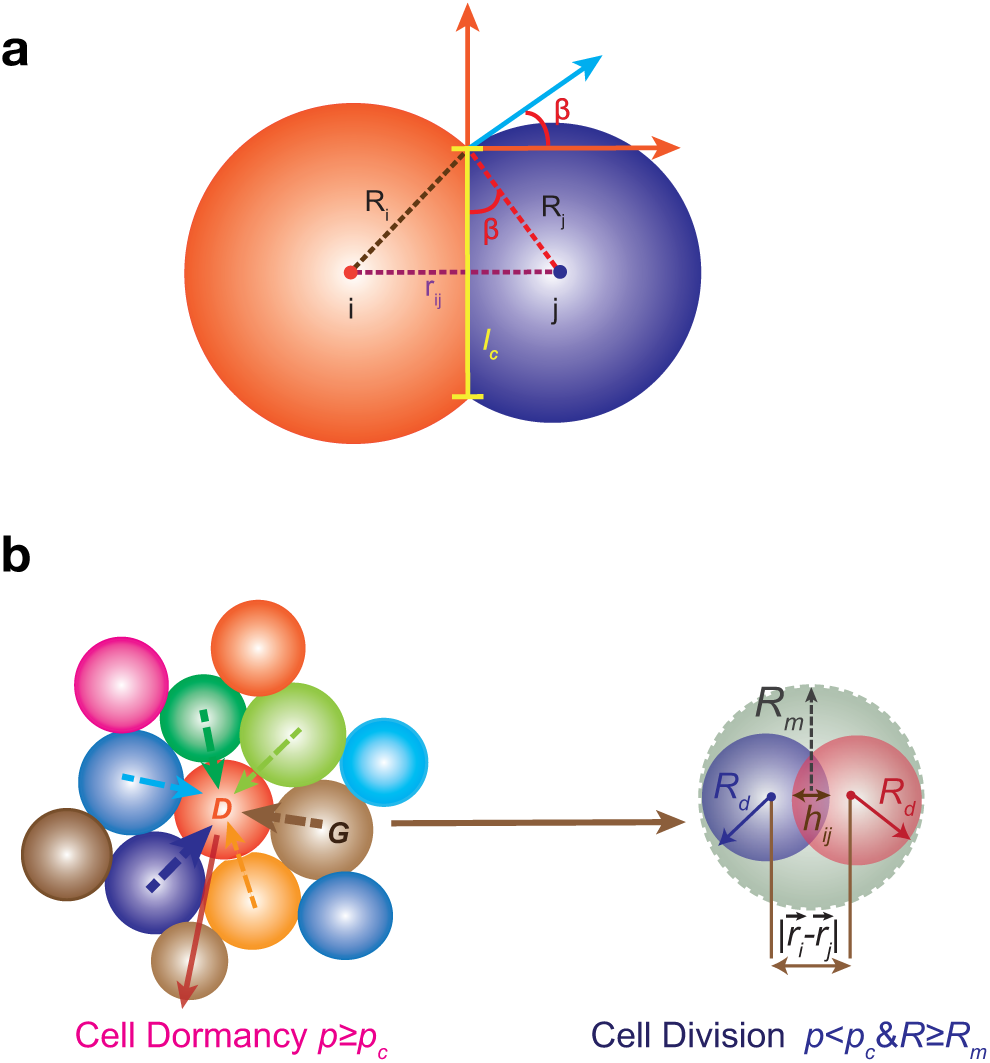
**a)** Cell-cell adhesion dictates the angle of contact between cells, *β*, and the length of contact, *l*_*c*_. **b)** Cell dormancy (left) and cell division (right). If the local pressure *p*_*i*_ that the *i*^*th*^ cell experiences (due to contacts with the neighboring cells) exceeds a specified critical pressure *p*_*c*_, it enters the dormant state (*D*). Otherwise, the cells undergo growth (G) until they reach the mitotic radius, *R*_*m*_. At that stage, the mother cell divides into two identical daughter cells with the same radius *R*_*d*_ in such a manner that the total volume upon cell division is conserved. A cell that is dormant at a given time can transit from that state at subsequent times. Cell center to center distance, 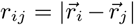, and cell overlap, *h*_*ij*_, are illustrated.

**FIG. 2.**
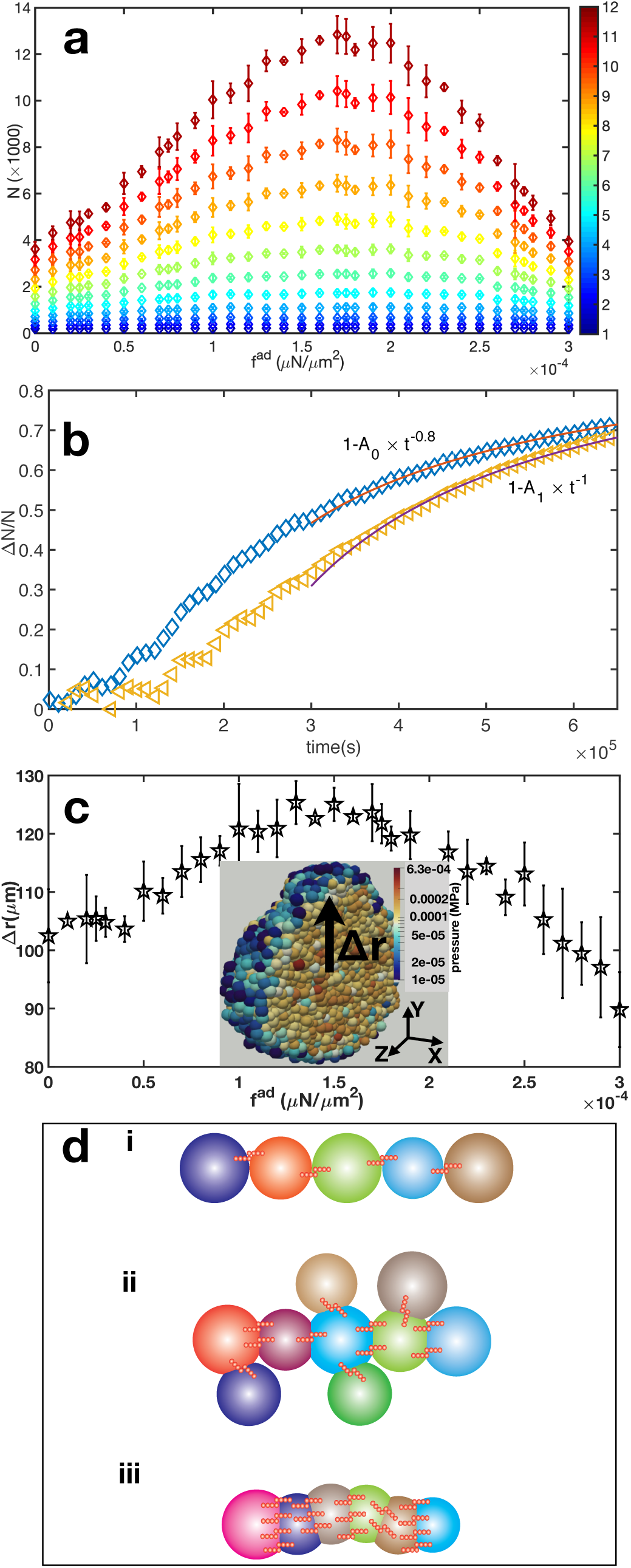
**a)** Proliferation capacity (PrC) measured as the total number of cells (N in units of 1000), at *t* = *τ*_*min*_ to 12*τ*_*min*_, at intervals of *τ*_*min*_(= 54, 000 sec), as a function of cell-cell adhesion strength (*f*^*ad*^) using *p*_*c*_ = 10^−4^MPa. Error bars here, and henceforth, are calculated from the standard deviation. The scale on the right gives *t* in units of *τ*_*min*_. **b)** Fractional change in *N* (defined in the text) between *f*^*ad*^ = 1.75 × 10^−4^*µ*N*/µ*m^2^ and *f*^*ad*^ = 0 (diamonds), between *f*^*ad*^ = 1.75 × 10^−4^ and *f*^*ad*^ = 3 × 10^−4^ (left triangles), as a function of time. Predicted power law behavior of Δ*N* (*t*)*/N* are plotted as lines. **c)** The dependence of the invasion distance (Δ*r*), at 7.5 days (= 12*τ*_*min*_), on *f*^*ad*^. Inset shows a cross section through a tumor spheroid and the distance Δ*r* from tumor center to periphery at *t* ∼ 12*τ*_*min*_ for *f*^*ad*^ = 1.5 × 10^−4^. Color indicates the pressure experienced by cells. **d)** Schematic for the three different regimes of cell-cell adhesion exhibiting differing PrCs. E-cadherin molecules are represented as short red bonds. These regimes in the simulations correspond to (i) *f*^*ad*^ ≤ 0.5 × 10^−4^ characterized by low cell-cell adhesion and low PrC. (ii) 1 × 10^−4^ ≤ *f*^*ad*^ ≤ 2 × 10^−4^ characterized by intermediate cell-cell adhesion and high PrC, and (iii) *f*^*ad*^ ≥ 2.5 × 10^−4^ with high cell-cell adhesion and low PrC.

**FIG. 3.**
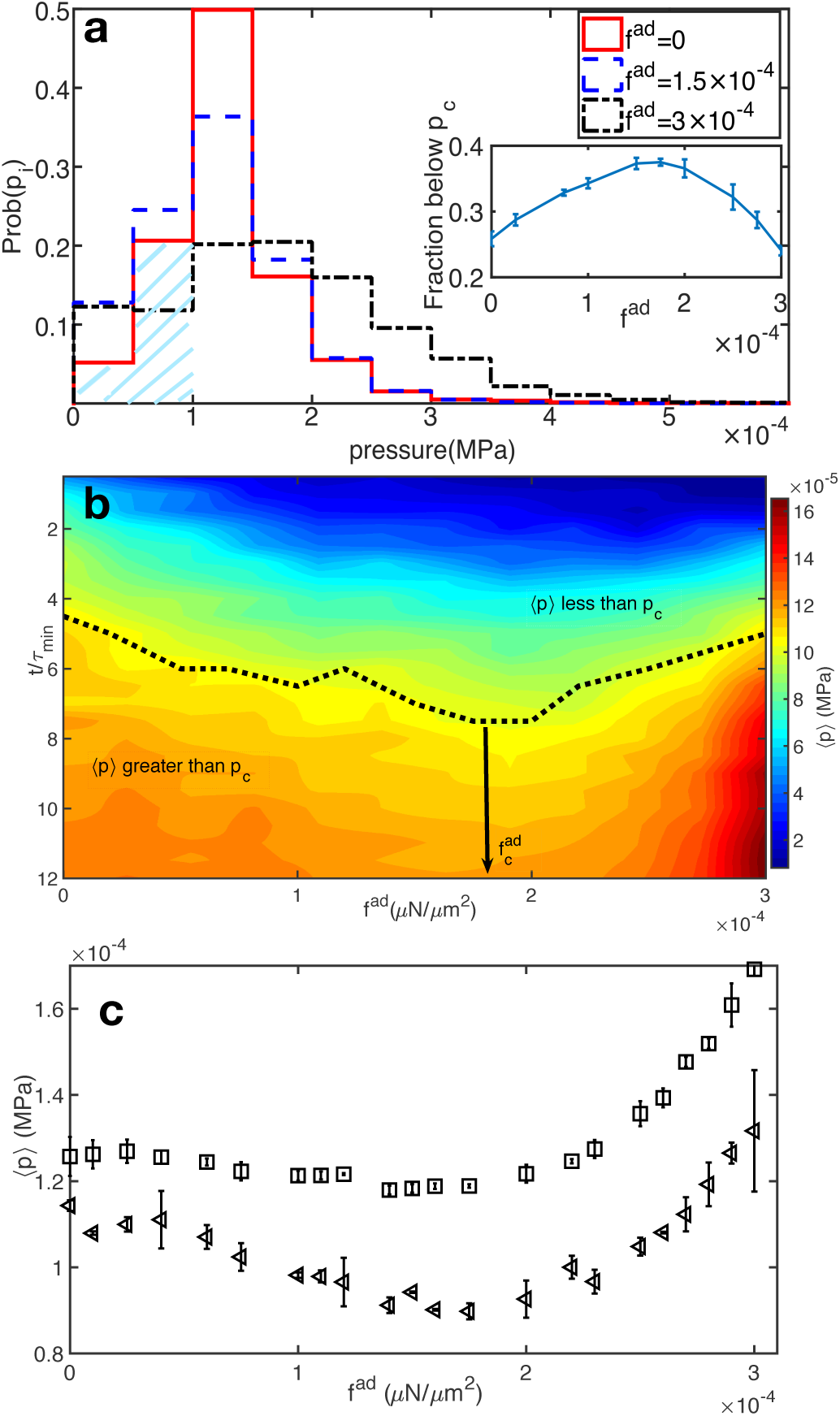
**a)** The probability distribution of pressure experienced by cells within a 3D tissue spheroid on day 7.5 for three different values of *f*^*ad*^. The shaded region represent pressure experienced by cells below *p*_*c*_ = 1 × 10^−4^MPa, at *f*^*ad*^ = 0. Fraction of cells (*F*_*C*_) with *p* < *p*_*c*_ is shown in the Inset. **b)** Phase plot of average pressure experienced by cells as a function of time and *f*^*ad*^. The scale for pressure is on the right. **c)** The average pressure ⟨*p*⟩ experienced by cells at *t* = 6*τ*_*min*_(left triangles; ∼ day 3.75 of growth) and *t* = 12*τ*_*min*_ (squares; ∼ day 7.5 of growth) for different values of *f*^*ad*^.

**FIG. 4.**
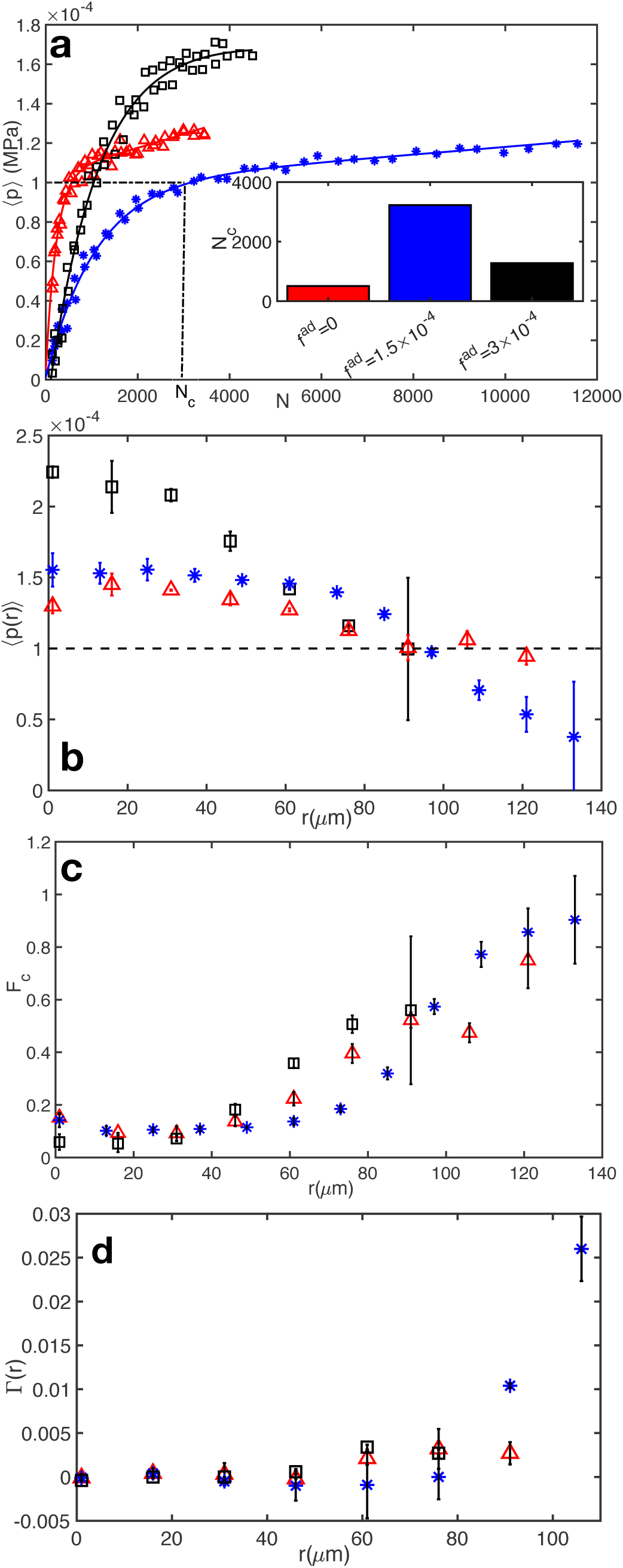
**a)**Average pressure experienced by cells, ⟨*p*⟩, as a function of the total number, *N*, of cells; ⟨*p*⟩ versus *N* at *f*^*ad*^ = 0 (red;triangle), *f*^*ad*^ = 1.5 × 10^−4^ (blue;asterisk) and *f*^*ad*^ = 3 × 10^−4^ (black;squares) are shown. The corresponding double exponential fits are, 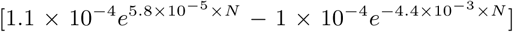 (red line, *f*^*ad*^ = 0), 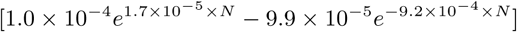 (blue line, *f*^*ad*^ = 1.5 × 10^−4^) and 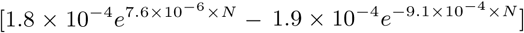 (black line, *f*^*ad*^ = 3 × 10^−4^). Inset shows *N*_*c*_, the number of cells at which ⟨*p*⟩ = *p*_*c*_. **b)** The average pressure experienced by cells at a distance *r* (*μ*m) from the spheroid center. The dashed line shows the critical pressure *p*_*c*_ = 10^−4^MPa. **c)** The fraction of cells with *p* < *p*_*c*_ at a distance *r*. **d)** The average cell proliferation rate at distance *r* from the center of the spheroid for *f*^*ad*^ = 0, *f*^*ad*^ = 1.5 ×10^−4^ and *f*^*ad*^ = 3 ×10^−4^. Colors and symbols corresponding to *f*^*ad*^ are the same for **a**-**d**. *t* is fixed at 650, 000*sec*(∼ 12*τ*_*min*_) for **b**-**d**.

**FIG. 5.**
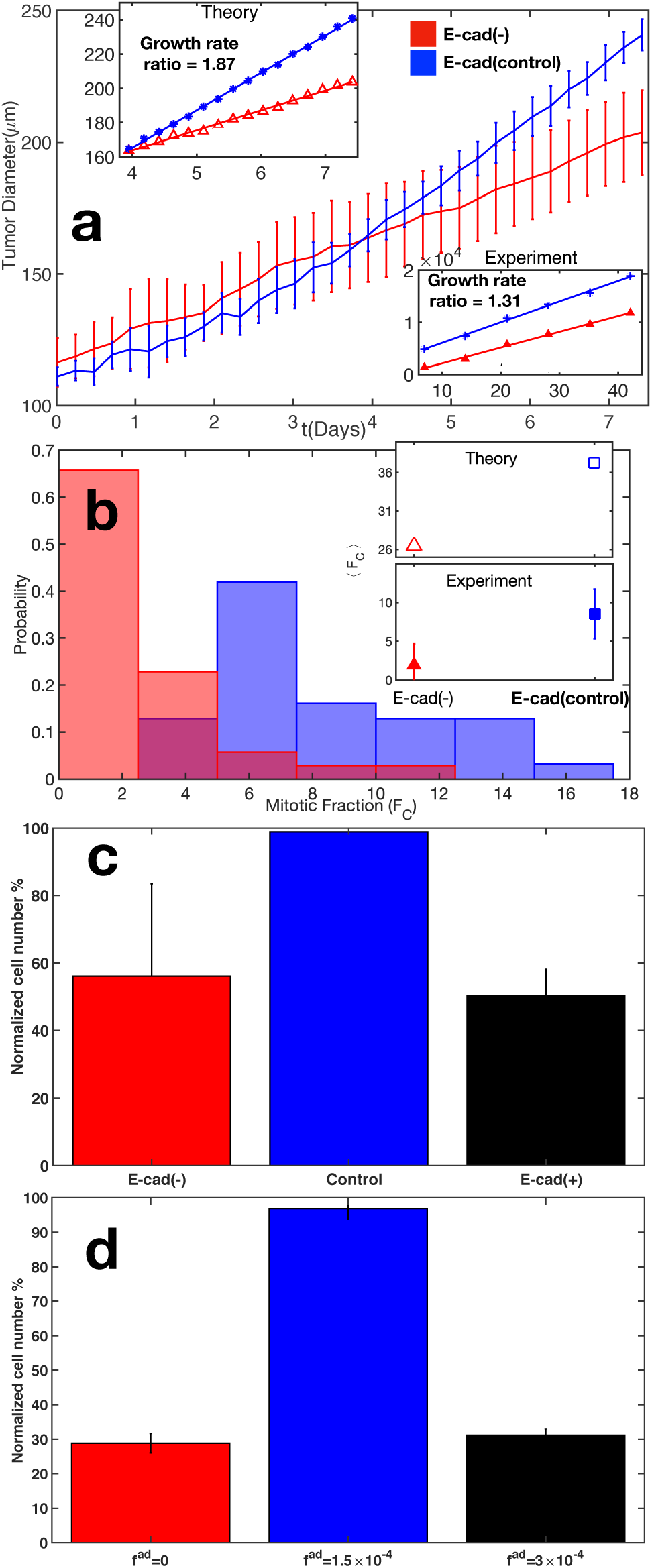
**a)** Main Panel) Kinetics of growth in the diameter of tumor spheroids composed of cells with low E-cadherin expression (red; *f*^*ad*^ = 0; E-cad(-)) and intermediate E-cadherin expression (blue; *f*^*ad*^ = 1.75 × 10^−4^; E-cad(control)) from simulations show enhanced tumor growth rate due to higher E-cadherin expression. The tumor diameter growth is linear at *t* > 4 days, independent of the E-cadherin expression level. However, the growth rate of the tumor colony with intermediate E-cadherin expression is larger (Top Inset; Theory) in agreement with experimental results. Growth rates of the longest tumor dimension in low E-cadherin expressing organoids compared to control organoids with normal E-cadherin expression is shown in the Bottom Inset (Data was extracted from Ref. [38]). **b)** Probability distribution of the mitotic fraction, *F*_*C*_, in cell colonies expressed as % for low E-cadherin tumor organoids (red) and normal E-cadherin expressing tumor organoids (blue) shows enhanced proliferation capacity when E-cadherin expression is increased. Data was extracted from Ref. [38]. (Top Inset; Theory) Mean mitotic fraction, ⟨*F*_*C*_⟩, for low E-cadherin tumor cells (E-cad(-)) and intermediate E-cadherin expressing tumor cells (E-cad (control)). (Bottom Inset; Experiment) Mean mitotic fraction in low E-cadherin expressing tumor organoids is lower than control organoids with normal E-cadherin expression [38]. Although there is paucity of experimental data it is encouraging that the currently available measurements are in line with our predictions. **c)** Experimental data for the number of cells normalized by maximum number (as %) at three levels of E-cadherin expression for primordial germ cells (PGC) from *Xenopus laevis* ≈ 1.5 days after ferilization [44]. Both overexpression and knockdown of E-cadherin leads to a decrease in PGC cell numbers. Control morpholino oligonucleotides (Contr MO) and uninjected show native E-cadherin expression while E-cad MO (E-cad(-)) and E-cad GFP (E-cad(+)) induces knockdown and overexpression of E-cadherin respectively. **d)** Simulation data for the number of cells normalized by maximum number (as %) at three levels of cell-cell adhesion strength (*t* = 650, 000*sec*(∼ 12*τ*_*min*_)). Values of *f*^*ad*^ (in units of *μN/μm*^2^) are shown in parenthesis, as a proxy for experimental E-cadherin expression levels.

**FIG. 6.**
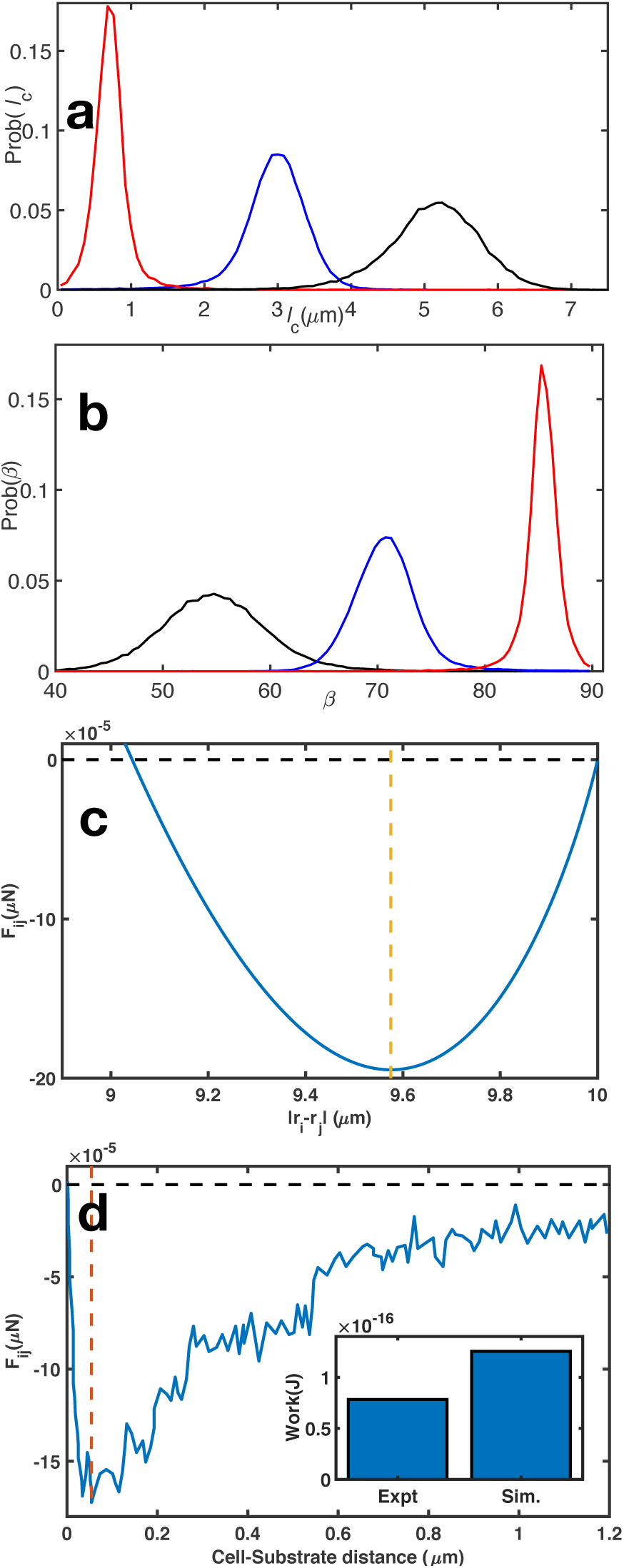
**a)** Probability distribution of contact lengths between cells for *f*^*ad*^ = 0 (red), *f*^*ad*^ = 1.5 × 10^−4^ (blue) and *f*^*ad*^ = 3 × 10^−4^ (black). **b)** Probability distribution of contact angles, *β* at varying values of *f*^*ad*^ with the color scheme as in **a). c)** Force on cell *i* due to *j, F*_*ij*_, for *R*_*i*_ = *R*_*j*_ = 5 *μm* using mean values of elastic modulus, poisson ratio, receptor and ligand concentration (see Table I in the SI). *F*_*ij*_ is plotted as a function of cell center-to-center distance |**r**_*i*_ −**r**_*j*_ |. We used *f*^*ad*^ = 1.75 × 10^−4^*μ*N*/μ*m^2^ to generate *F*_*ij*_. **d)** Force-distance data extracted from SCFS experiment [44]. Inset shows the work required to separate cell and E-cadherin functionalized substrate in SCFS experiment and two cells in theory, respectively. Minimum force values are indicated by vertical dashed lines. See Appendix B for further details.

Dynamics of cell-cell contact leads to spatiotemporal fluctuations in the local forces experienced by a cell (and vice versa), through which we implement a mechanical feedback on growth and proliferation. Depending on the local forces, a cell can either be in the dormant (*D*) or in the growth (*G*) phase (see Fig 1b). The effect of the local cell microenvironment on proliferation, a collective cell effect, is taken into account through the pressure experienced by the cell (*p*_*i*_; see Appendix A). We refer to *p*_*i*_ as pressure since it has the same dimensions. However, it is not as rigorous a definition of pressure as the thermo-dynamic conjugate of the volume. Essentially, *p*_*i*_, models the mechanical sensitivity of cell proliferation to the local environment. If *p*_*i*_ exceeds *p*_*c*_ (a pre-assigned value of the critical pressure), the cell enters dormancy (D), and can no longer grow or divide. However, if *p*_*i*_ < *p*_*c*_, the cell can continue to grow in size until it reaches the mitotic radius *R*_*m*_, the size at which cell division occurs.

### Proliferation Depends Non-monotonically on *f* ^*ad*^

Our major finding is summarized in Fig. 2, which shows that tumor proliferation and a measure of invasiveness (Δ*r*(*t* = 7.5 days)), exhibit a non-monotonic dependence on *f*^*ad*^. Both quantities increase from small values as *f*^*ad*^ increases, attain a maximum at 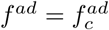, and decrease as *f*^*ad*^ exceeds 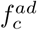. In the simulations, we began with 100 cells at *t* = 0, and let the cells evolve as determined by the equations of motion, and the rules governing birth and apoptosis (see Appendix A). We performed simulations at different values of *f*^*ad*^ until ∼ 7.5 days (= 12*τ*_*min*_, where *τ*_*min*_ is the average cell division time), sufficient to account for multiple cell division cycles. The total number of cells (*N*) as a function of *f*^*ad*^ at various times in the range of 1*τ*_*min*_ to 12*τ*_*min*_ (with colors for time *t* in units of *τ*_*min*_) are shown in Fig. 2a.

On increasing *f*^*ad*^ from 0 (no E-cadherin expression) to 1.75 × 10^−4^*µ*N*/µ*m^2^ (intermediate E-cadherin expression), the total number of cells, *N*, at *t* = 12*τ*_*min*_(∼7.5 days; dark red in Fig. 2a) increases substantially. When *f*^*ad*^ exceeds 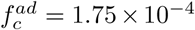, the proliferation capacity (PrC) is down-regulated (Fig. 2a). While *N* = 12, 000 cells on day 7.5 at *f*^*ad*^ = 1.75 × 10^−4^*µ*N*/µ*m^2^, for higher values (3 × 10^−4^*µ*N*/µ*m^2^), the tumor consists of only 4, 000 at the same *t*. The surprising non-monotonic dependence of cell numbers, *N*, on *f*^*ad*^, is qualitatively consistent with some recent experiments [36, 38, 44], as we discuss below. The non-monotonic proliferation behavior becomes pronounced beginning at *t* = 5*τ*_*min*_ (see Fig. 2a). The fractional change in the number of cells between 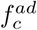 and another value of 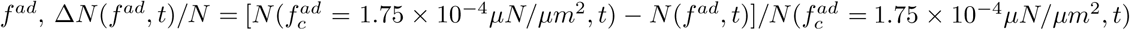 as a function of time, quantifies the asymmetry in the proliferation capacity due to adhesion strength. As shown in Fig. 2b, the parameter Δ*N* (*t*)*/N*, exhibits non-linear behavior. From fits of *N*, as a function of time, the proliferation asymmetry parameter is expected to evolve in time as 1 − *A*_0_ × *t*^−0.8^ for *f*^*ad*^ = 0 and 1 − *A*_1_ × *t*^−1^ for *f*^*ad*^ = 3 × 10^−4^*µN/µm*^2^, where *A*_0_ and *A*_1_ are constants (see Appendix A Fig. 7a for the power law fits).

**FIG. 7.**
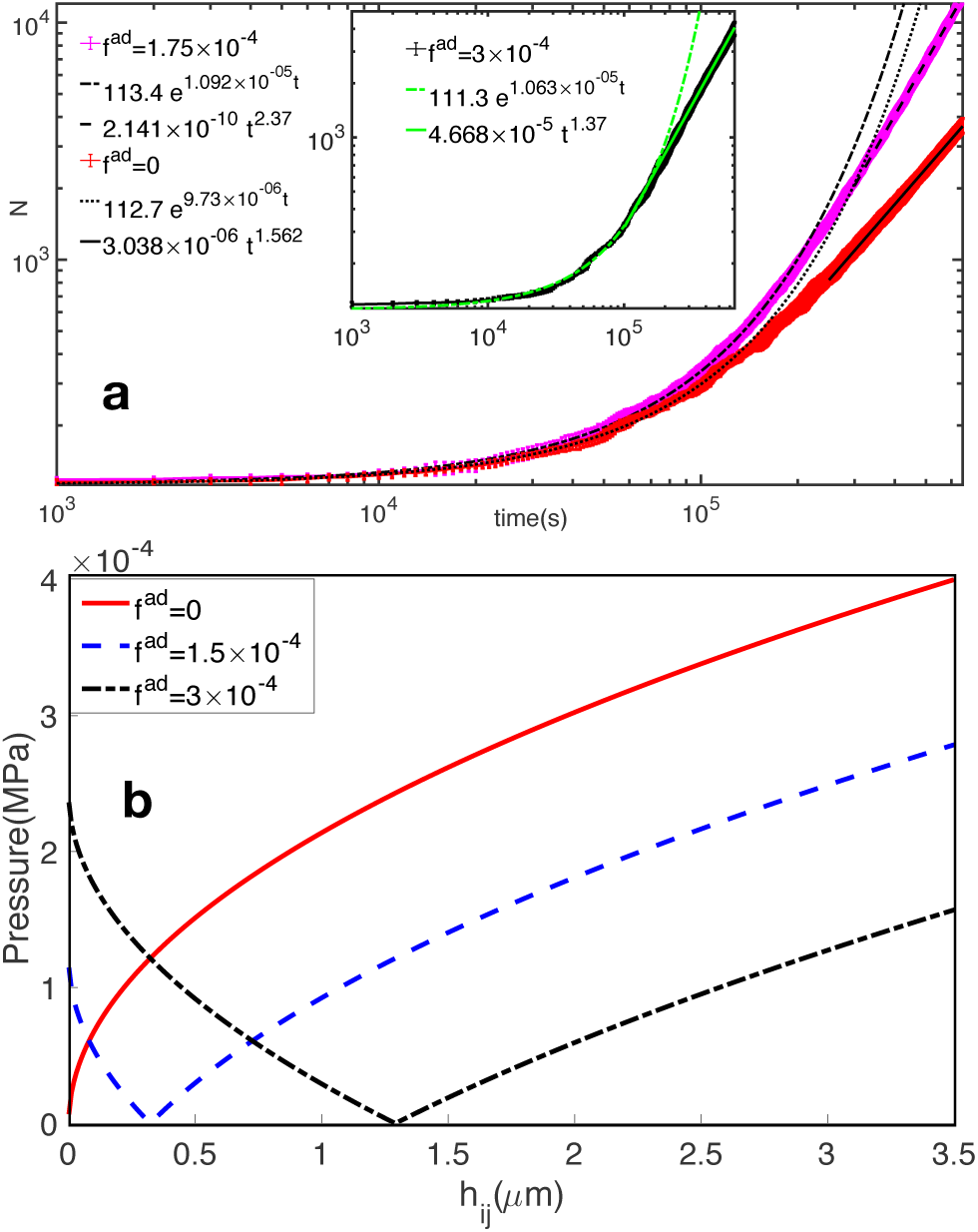
**a)** Number of cells, *N* (*t*), over 7.5 days of growth. Initial exponential growth followed by power law growth behavior is seen for three different *f*^*ad*^ values. The onset of power-law growth in *N* occurs between *t* = 10^5^ − 2 × 10^5^ secs. Power law exponent depends on *f*^*ad*^. **b)** Pressure, *p*_*i*_, experienced by a cell interacting with another cell as a function of overlap distance (*h*_*ij*_) for different values of *f*^*ad*^. *F*_*ij*_ is calculated for *R*_*i*_ = *R*_*j*_ = 4 *μm* using mean values of elastic modulus, poisson ratio, receptor and ligand concentration (see Table I).

### Invasion Distance Mirrors Tumor Proliferation Behavior

The invasion or spreading distance, Δ*r*(*t*) (shown in Fig. 2c), measurable experimentally using imaging methods [45], is the average distance between center of mass of the tumor spheroid and the cells at tumor periphery, 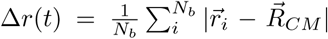. Here, the summation is over *N*_*b*_, the number of cells at the tumor periphery at positions 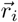 and 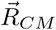 is the tumor center of mass 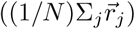. In accord with increased proliferation shown in Fig. 2a, Δ*r*(*t* = 12*τ*_*min*_) is also enhanced at intermediate values of *f*^*ad*^ (Fig. 2c). The uptick in invasiveness from low to intermediate values of *f*^*ad*^ is fundamentally different from what is expected in the canonical picture, where increasing cell-cell adhesion suppresses invasiveness and metastatic dissemination of cancer cells [29, 46]. In contrast, tumor invasiveness as a function of increasing adhesion or stickiness between cells (as tracked by Δ*r*(*t* = 12*τ*_*min*_)) initially increases and reaches a maximum, followed by a crossover to a regime of decreasing invasiveness at higher adhesion strengths. We note that the decreased invasive behavior at 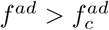 is in agreement with the canonical picture, where enhanced E-cadherin expression results in tumor suppression. Schematic summary of the results is presented in Fig. 2d.

The inset in Fig. 2c shows the highly heterogenous spatial distribution of intercellular pressure (snapshot at *t* = 12*τ*_*min*_), marked by elevated pressure at the core and decreasing as one approaches the tissue periphery. As cell rearrangement and birth-death events give rise to local cell density fluctuations, the cell pressure is a highly dynamic quantity, see videos (Supplementary Movies 1-3; Appendix A Figs. 8a-c) for illustration of pressure dynamics during the growth of the cell collective. Spatial distribution of pressure is important to understanding the non-monotonic proliferation behavior, as we discuss in more detail below.

**FIG. 8.**
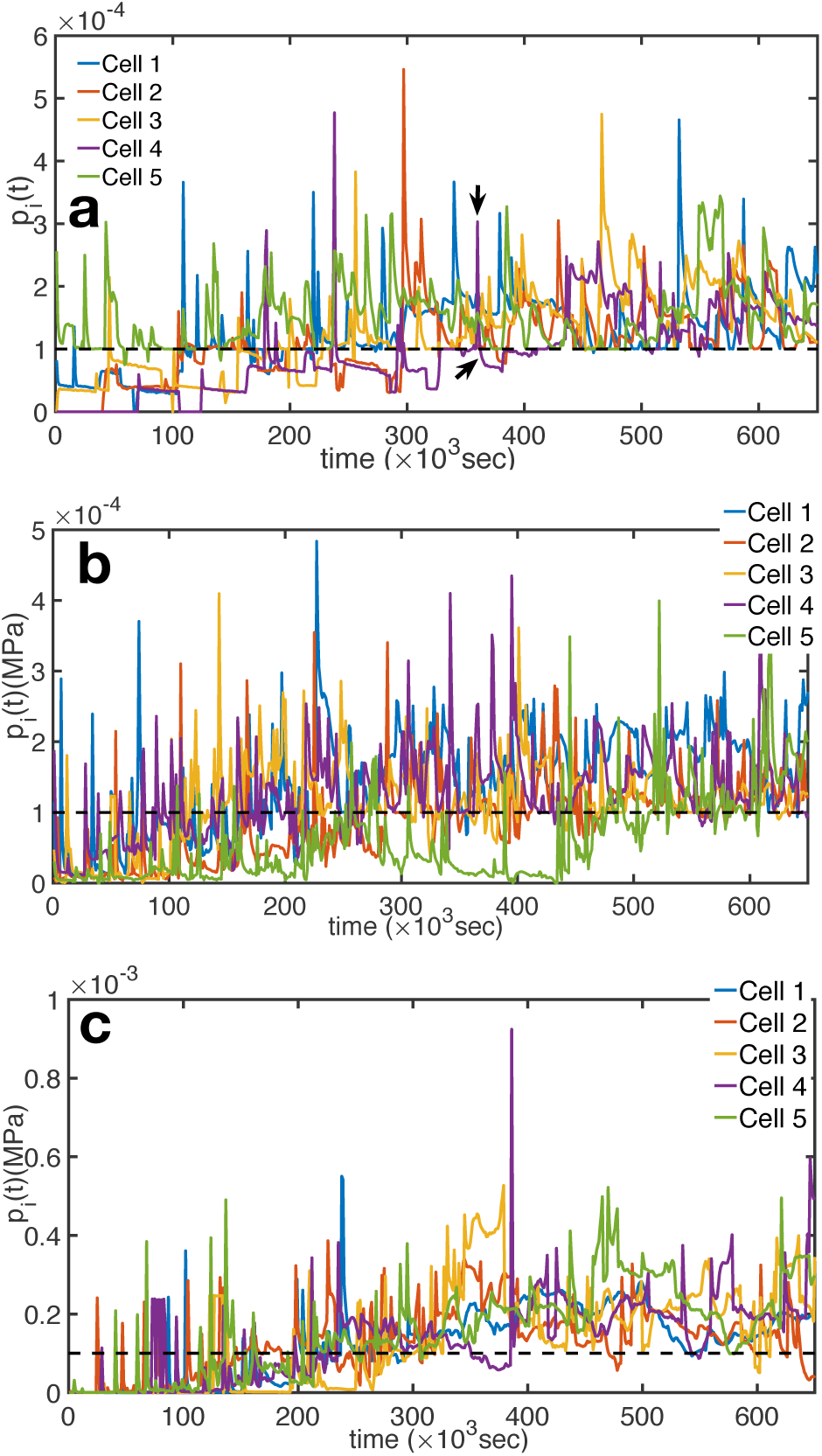
**a)** Pressure experienced by individual cells as a function of time at *f*^*ad*^ = 0, **b)** *f*^*ad*^ = 1.5 × 10^−4^ and **c)** *f*^*ad*^ = 3 × 10^−4^. Dashed black lines indicate the critical pressure, *p*_*c*_. Black arrows in **a)** highlight fluctuations in *p*_*i*_ above and below *p*_*c*_.

### Fraction of Dormant Cells Determine Non-monotonic Proliferation

To understand the physical factors underlying the non-monotonic proliferation behavior shown in Fig. 2a, we searched for the growth control mechanism. We found that the pressure experienced by a cell as a function of cell packing in the 3D spheroid (Appendix A Fig. 7b) plays an essential role in the observed results. For *f*^*ad*^ > 0, a minimum in pressure is observed at non-zero cell-cell over-lap distance, *h*_*ij*_ (Appendix A Fig. 7b). For instance, at *f*^*ad*^ = 1.5 × 10^−4^*µ*N*/µ*m^2^ and 3 × 10^−4^*µ*N*/µ*m^2^, the pressure (*p*_*i*_) experienced by cells is zero at *h*_*ij*_ ∼ 0.4*µ*m and 1.3*µ*m respectively. At this minimum pressure, *p*_*i*_ → 0, the proliferation capacity (PrC) of the cells is maximized because cells are readily outside the dormant regime, *p*_*i*_*/p*_*c*_ < 1. Due to the relationship between cell-cell overlap (*h*_*ij*_) and center-to-center distance (|**r**_*i*_ − **r**_*j*_| = *R*_*i*_ + *R*_*j*_ − *h*_*ij*_), our conclusions regarding *h*_*ij*_ can equivalently be discussed in terms of the cell-cell contact length (*l*_*c*_), the angle *β* (see Fig. 1) and cell-cell internuclear distance, |*r*_*i*_ − *r*_*j*_|.

Fig. 3a shows the probability distribution of pressure at *t* = 7.5 days at three representative values of *f*^*ad*^, corresponding to low, intermediate and high adhesion strengths. The crucial feature that gives rise to the non-monotonic proliferation and invasion (Figs. 2a -2c) is the nonlinear behavior of the pressure distribution below *p*_*c*_ = 1 × 10^−4^MPa (see shaded portion in Fig. 3a, quantified in the Inset) as *f*^*ad*^ is changed. In the inset of Fig. 3a, the fraction of cells in the proliferating regime shows a biphasic behavior with a peak at 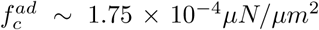. The fraction of cells in the growth phase (*p*_*i*_ < *p*_*c*_) peaks at ≈ 38%, between 1 × 10^−4^ *µ*N*/µ*m^2^ < *f*^*ad*^ < 2 × 10^−4^ *µ*N*/µ*m^2^. For both lower and higher cell-cell adhesion strengths, the fraction of cells in the growth phase is at or below 25% on day 7.5 of tissue growth. We present the phase diagram of the average pressure 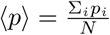, color map) on cells as a function of *f*^*ad*^ and time (in units of *τ*_*min*_) in Fig. 3b. The dotted line marks the boundary ⟨*p*⟩ = *p*_*c*_, between the regimes where cells on average grow and divide (*p*_*i*_ < *p*_*c*_), and the opposite limit where they are dormant. At a fixed value of *t*, the regime between 1 × 10^−4^ ≤ *f*^*ad*^ ≤ 2 × 10^−4^ shows a marked dip in the average pressure experienced by cells. The low pressure regime, ⟨*p*⟩ < *p*_*c*_, is particularly pronounced between 2*τ*_*min*_ ≤ *t* ≤ 7*τ*_*min*_. In Fig. 3c, the average pressure at *t* = 6*τ*_*min*_ and 12*τ*_*min*_, as a function of *f*^*ad*^, are shown for illustration purposes. Minimum in ⟨*p*⟩ between *f*^*ad*^ ≈ 1.5 − 2 × 10^−4^ *µ*N*/µ*m^2^ is seen. In our pressure dependent model of contact inhibition, there is a close relationship between proliferation capacity (PrC) and the local pressure on a cell. As the number of cells experiencing pressure below critical pressure increases, the PrC of the tissue is enhanced. It is less obvious why the average cell pressure acquires a minimum value at 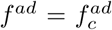 (Fig. 3b-c). We provide a plausible theoretical explanation below.

### Pressure Gradient Drives Biphasic Proliferation Behavior

The finding in Fig. 2a could be understood using the following physical picture. PrC is determined by the number of cells with pressures less than *p*_*c*_, the critical value. If the pressure on a cell exceeds *p*_*c*_, it becomes dormant, thus losing the ability to divide and grow. The average pressure that a cell experiences depends on the magnitude of the net adhesive force. At low values of *f*^*ad*^ (incrementing from *f*^*ad*^ = 0) the cell pressure decreases as the cells overlap with each other because they are deformable (|*r*_*i*_ − *r*_*j*_| < *R*_*i*_ + *R*_*j*_). This is similar to the pressure in real gases (Van der Waals picture) in which the inter-particle attraction leads to a decrease in the average pressure. As a result, in a certain range of *f*^*ad*^, we expect the number of cells capable of dividing should increase, causing enhanced proliferation. At very high values of *f*^*ad*^, however, the attraction becomes so strong that the number of nearest neighbors of a given cell increases (Appendix C Figs. 9a-b). This leads to an overall increase in pressure (jamming effect), resulting in a decrease in the number of cells that can proliferate. From these arguments it follows that *N* (*t*) should increase (decrease) at low (high) *f*^*ad*^ values with a maximum at an intermediate *f*^*ad*^. The physical picture given above can be used to construct an approximate theory for the finding that cell proliferation reaches a maximum at 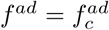 (Fig. 2a). The total average pressure (*p*_*t*_) that a cell experiences is given by 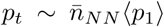 where 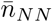 is the mean number of nearest neighbors, and ⟨*p*_1_⟩ is the average pressure a single neighboring cell exerts on a given cell. We appeal to simulations to estimate the dependence of 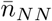 on *f*^*ad*^ (see Appendix C Figs. 9a-b). For any cell *i*, the nearest neighbors are defined as those cells with non-zero overlap (i.e. *h*_*ij*_ > 0). To obtain ⟨*p*_1_⟩, we expand around *h*_0_, the cell-cell over-lap value where both the attractive and repulsive interaction terms are equal 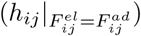, corresponding to the overlap distance at which *p* = 0 (Appendix A Fig. 7b). Thus, by Taylor expansion to first order, ⟨*p*_1_⟩ can be written as 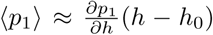. Here, 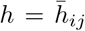, is the mean cell-cell overlap, which depends on *f*^*ad*^ (see Appendix C Fig. 10). We note that the variation in *h*_0_ with respect to *R*_*i*_ and *R*_*j*_, as well as other cell-cell interaction parameters is small compared to cell size. Estimating the dependence of *h* − *h*_0_ on adhesion strength (see Appendix C Fig. 11), an approximate linear trend is observed. At higher adhesion strengths, cells find it increasingly difficult to rearrange themselves and pack in such a way that intercellular distances are optimal for proliferation.

**FIG. 9.**
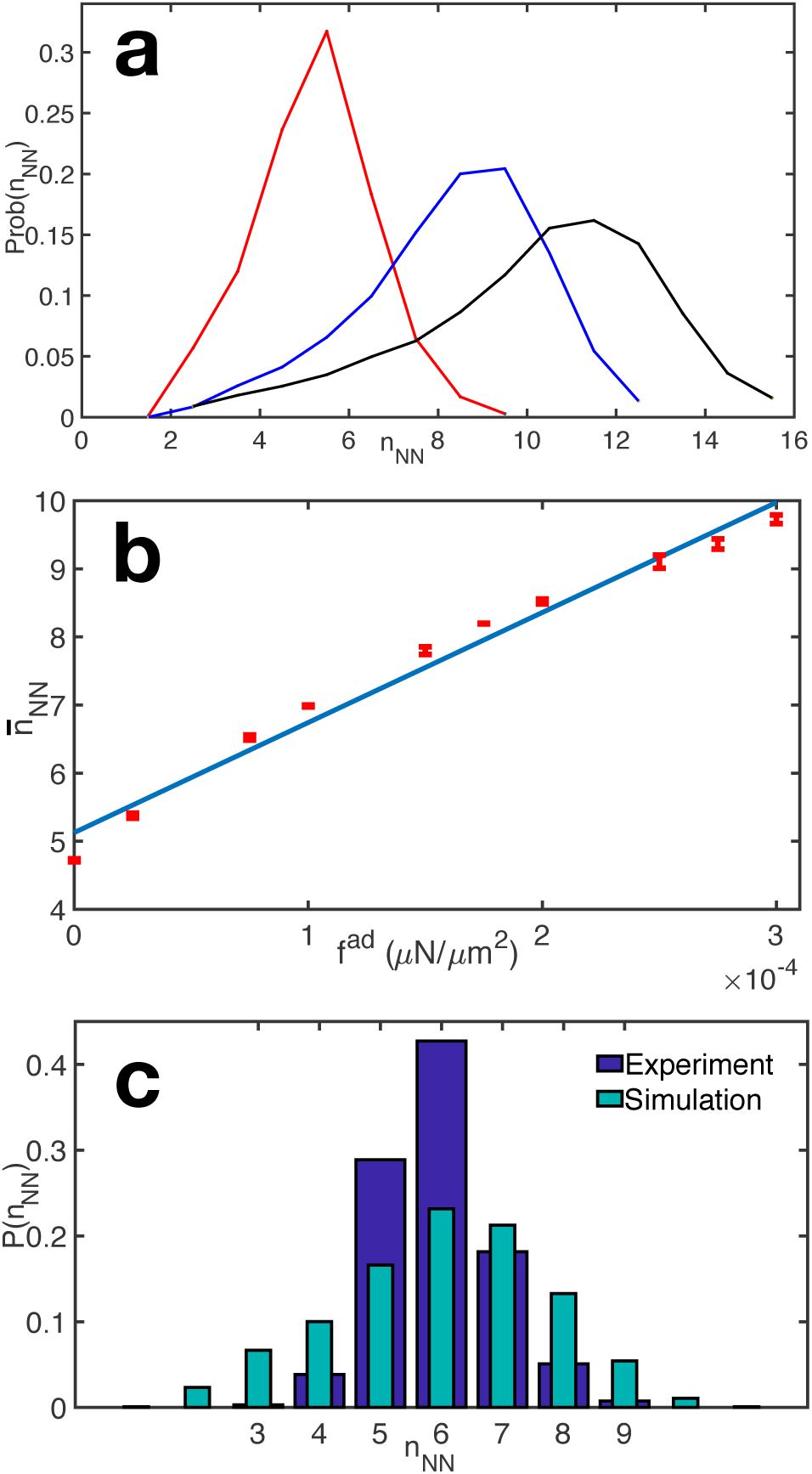
**a)** Graph showing the probability distribution of the number of nearest neighbors (*n*_*NN*_) on day 7.5 of tumor growth. Red, blue and black curves are for *f*^*ad*^ = 0, 1.5 × 10^−4^*μ*N*/μ*m^2^, and 3 × 10^−4^*μ*N*/μ*m^2^, respectively. **b)**Average number of nearest neighbors, 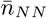, as a function of *f*^*ad*^. Linear fit shows 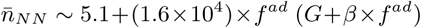. Error bars represent the standard deviation. **c)** Comparison of the probability distribution of the number of nearest neighbors (*n*_*NN*_) between experiments and simulations. Experimental data is for Xenopus tail epidermis (n=1,051 cells) [77]. Simulation data is for *f*^*ad*^ = 5 ×10^−5^ *μ*N*/μ*m^2^. Data in (a) - (c) are from 3 independent simulation runs.

**FIG. 10.**
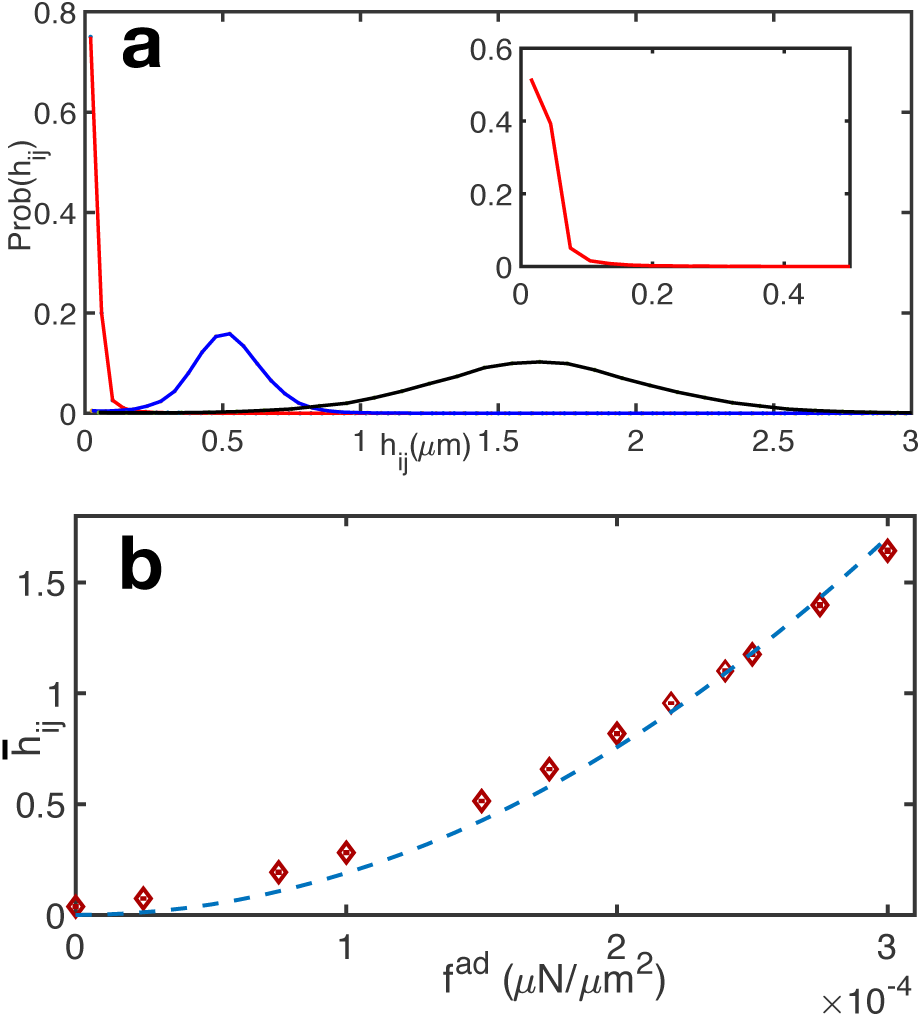
**a)** Probability distribution of the overlap (*h*_*ij*_) of cells on day 7.5 of tumor growth for three different adhesion strengths. Red, blue and black curves are for *f*^*ad*^ = 0, 1.5 × 10^−4^*µN/µm*^2^, and 3 × 10^−4^*µN/µm*^2^, respectively. **b)** Average interpenetration distance has a quadratic dependence on adhesion strength - 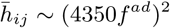. Data obtained are from 3 independent simulation runs. Error bars represent the standard deviation.

**FIG. 11.**
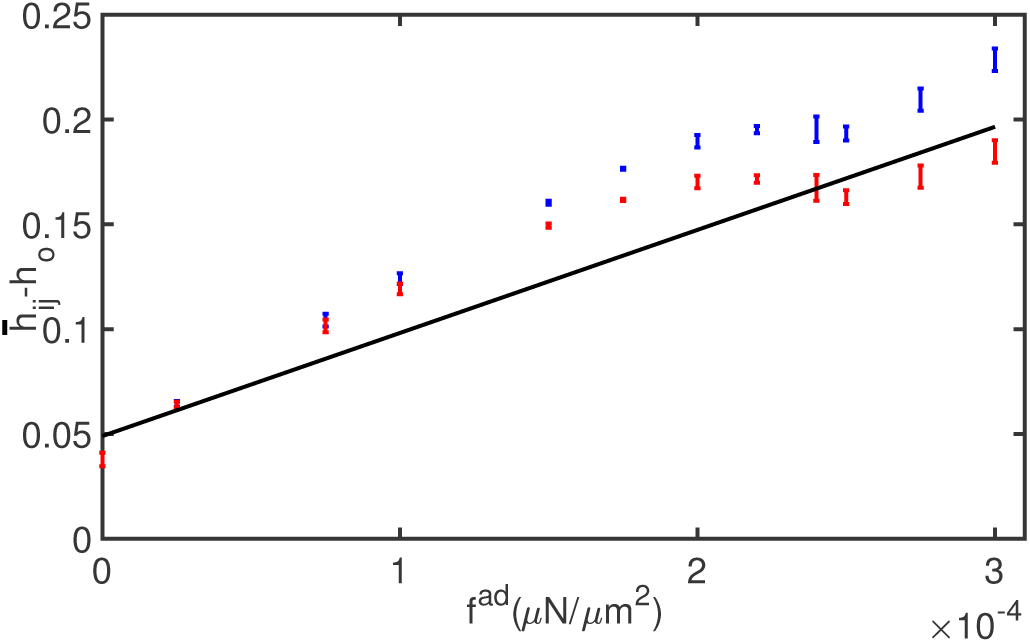
Deviation of the average 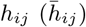 from *h*_0_ (cell interpenetration distance with minimum possible pressure) for differing adhesion strength showing approximate linear dependence 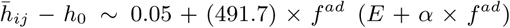. The value of *h*_0_ is determined by the radii of any two interacting cells. Two sets of points are shown, blue bars are for average radii of *R*_*i*_ = *R*_*j*_ = 3.9*µm* and the reds are for *R*_*i*_ = *R*_*j*_ = 4*µm.*

In order to calculate the pressure gradient with respect to cell-cell overlap, 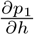, we use

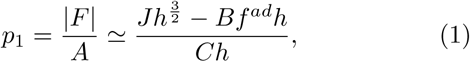

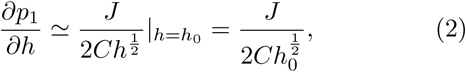

to separate the dependence on cell-cell overlap and adhesion strength. In Eqs. 1 and 2 *J, B* and *C* are independent of both *h* and *f*^*ad*^ and can be obtained from Appendix A Eqs. A2 and A3, and the definition of *A*_*ij*_. The resulting expressions are: 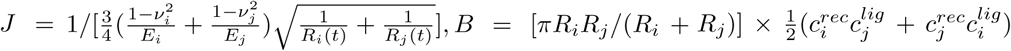, and *C* = *πR*_*i*_*R*_*j*_/(*R*_*i*_ + *R*_*i*_). On equating the repulsive and attractive interaction terms, we obtain *h*_0_ ≈ *K*(*f*^*ad*^)^2^, implying 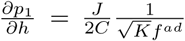, where, *K* = (*B/J*)^2^. Thus, the total pressure (*p*_*t*_) experienced by a cell is 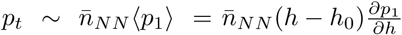,

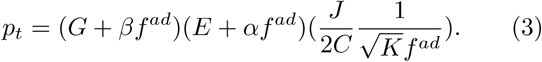

The mean number of near-neighbors 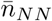 increases with *f*^*ad*^ and to a first approximation can be written as *G* + *βf*^*ad*^ (see Appendix C Fig. 9a-b; *G, β* are constants obtained from fitting simulation data). Similarly, the deviation of the cell-cell overlap from *h*_0_ is approximately *E* + *αf*^*ad*^ (see Appendix C Fig. 11; *E, α* are constants). Notice that Eq. 3 can be written as, 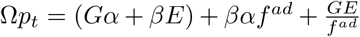, where 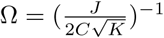. In this form, the second term depends linearly on *f*^*ad*^ and the third is inversely proportional to *f*^*ad*^. As described in the physical arguments, enhancement in proliferation is maximized if *p*_*t*_ is as small as possible. The minimum in the total pressure experienced is given by the solution to 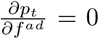. Therefore, the predicted optimal cell-cell adhesion strength is 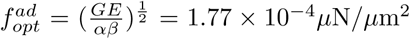. This is in excellent agreement with the simulation results 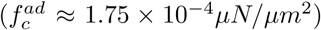 for the peak in the proliferation behavior (*N* (*t* = 7.5 days) in Fig. 2a). More importantly, the arguments leading to Eq. 3 show that the variations in the average pressure as a function of *f*^*ad*^ drives proliferation.

### Cell Packing and Spatial Proliferation Patterns Are Dictated by Cell-Cell Adhesion

Mechanosensitivity i.e. specialized response to mechanical stimulation is common to cells in many organisms. Exposure to stresses in living tissues can module physiological processes from molecular, to cellular and systemic levels [47]. Here, we quantify the spatio-temporal behavior of the intercellular pressure, the parameter that encodes the mechanical sensitivity of cell proliferation to the local environment, and delineate its emergent properties in a growing cell collective. The average pressure, ⟨*p*⟩, experienced by cells as a function of the total number of cells at varying *f*^*ad*^ is shown in Fig 4a. *N*_*c*_, defined as the number of cells at which ⟨*p*⟩ = *p*_*c*_, exhibits a biphasic behavior, supporting the maximum number of cells at an intermediate *f*^*ad*^. This provides further evidence that in a growing collection of cells, at intermediate *f*^*ad*^, cells rearrange and pack effectively in such a manner that the average pressure is minimized. For all three adhesion strengths considered, an initial regime where pressure rises rapidly is followed by a more gradual increase in pressure, coinciding with the exponential to power law crossover in the growth in the number of cells (see Fig. 7a in Appendix A). ⟨*p*⟩ as a function of *N* are well fit by double exponential functions. We propose that the intercellular pressure in 3D cell collectives should exhibit a double exponential dependence on *N*. However, the precise nature of the intercellular pressure depends on the details of the interaction between cells.

With recent advances in experimental techniques [48, 49], it is now possible to map spatial variations in intercellular forces within 3D tissues. Hence, we study how the spatial distribution of pressure and proliferation is influenced by cell-cell adhesion strength. We find that the cells at the tumor center experience higher pressures, above the critical pressure *p*_*c*_ (see Fig. 4b), independent of *f*^*ad*^. In contrast, the average pressure experienced by cells close to the tumor periphery is below the critical value *p*_*c*_. The pressure decreases as a function of distance *r* from the tumor center, with the lowest average pressure observed at the intermediate value of *f*^*ad*^ = 1.5 × 10^−4^ as one approaches the tumor periphery. We calculate the average pressure dependence on *r* using,

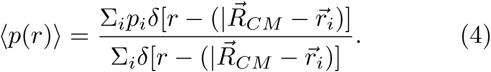

Due to the high pressure experienced by cells near the tumor center, a low fraction (*F*_*c*_ < 0.2) of cells are in growth phase at small *r* < 50*µm* while the majority of cells can grow at large *r* (see Fig. 4c). A rapid increase in *F*_*c*_ is observed approaching the tumor periphery for the intermediate value of *f*^*ad*^ = 1.5 × 10^−4^ (see the blue asterisks in Fig. 4c). To understand the rapid tumor invasion at intermediate value of *f*^*ad*^, we calculated the average cell proliferation rate, G(*r*), at distance *r* from the tumor center, 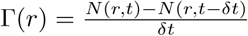. Here, *N* (*r, t*) is the number of cells at time *t* = 650, 000*s* and *δt* = 5000*s* is the time interval. The average is over polar and azimuthal angles, for all cells between *r* to *r* + *δr*. Closer to the tumor center, at low *r*, Γ(*r*) ∼ 0 indicating no proliferative activity. However, for larger values of *r*, proliferation rate rapidly increases approaching the periphery. We found that the cell proliferation rate is similar for different *f*^*ad*^ at small *r*, while a much higher proliferation rate is observed for the intermediate value of *f*^*ad*^ at larger *r* (see Fig. 4d). This spatial proliferation profile is in agreement with experimental results, where increased mechanical stress is correlated with lack of proliferation within the spheroid core, albeit in the context of externally applied stress [48]. We show, however, that even in the absence of an external applied stress, intercellular interactions give rise heterogeneity in the spatial distribution of mechanical stresses.

## DISCUSSION

### Internal Pressure Provides Feedback In E-cadherin’s Role In Tumor Dynamics

We have shown that the growth and invasiveness of the tumor spheroid changes non-monotonically with *f*^*ad*^, exhibiting a maximum at an intermediate value of the inter-cell adhesion strength. The mechanism for this unexpected finding is related to a collective effect that alters the pressure on a cell due to its neighbors. Thus, internal pressure may be viewed as providing a feedback mechanism in tumor growth and inhibition. The optimal value of 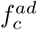 at which cell proliferation is a maximum at *t* >> *τ*_*min*_, is due to *p*_*c*_, the critical pressure above which a cell enters dormancy. Taken together these results show that the observed non-monotonic behavior is due to an interplay of *f*^*ad*^ and pressure, which serves as a feedback in enhancing or suppressing tumor growth. We note that the main conclusion of our model, on the non-monotonic proliferation behavior with a maximum at 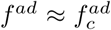, is independent of the exact value of *p*_*c*_ (see SI Fig. S1), alternative definitions of pressure experienced by the cells as well as the cell-cell interaction (see SI Figs. S2-S3; see SI Section I and II for more details).

The growth mechanism leading to non-monotonic proliferation at times exceeding a few cell division cycles is determined by the fraction of cells, *F*_*C*_, with pressure less than *p*_*c*_. The growth rate, and hence *F*_*C*_, depends on both *p*_*c*_ as well as *f*^*ad*^. This picture, arising in our simulations, is very similar to the mechanical feedback as a control mechanism for tissue growth proposed by Shraiman [15, 21]. In his formulation, the tissue is densely packed (perhaps confluent) so that cellular rearrangements does not occur readily, and the tissue could be treated as an elastic sheet that resists shear. For this case, Shraiman produced a theory for uniform tissue growth by proposing that mechanical stresses serves as a feedback mechanism. In our case, large scale cell rearrangements are possible as a cell or group of cells could go in and out of dormancy, determined by the *p*_*i*_(*t*)*/p*_*c*_. Despite the differences between the two studies, the idea that pressure could serve as a regulatory mechanism of growth, which in our case leads to non-monotonic dependence of proliferation on cell-cell adhesive interaction strength, could be a general feature of mechanical feedback[15].

### Cell-matrix Interactions

Biphasic cell migration controlled by adhesion to the extracellular matrix has been previously established. In pioneering studies, Lauffenburger and coworkers [50, 51] showed using theory and experiments that speed of single cell migration is biphasic, achieving an optimal value at an intermediate strength of cell-substratum interactions. Similarly, invasion of melanoma cells into a matrix was also found to be biphasic [52], increasing at small collagen concentrations and reaching a peak at an intermediate value. At much higher values of the collagen concentration, the invasion into the matrix decreases giving rise to the biphasic dependence. A direct link between the survival of genetically induced glioma-bearing mice and the expression level of CD44, which is a cell surface marker, has recently been reported [53]. The authors showed that the survival depends in a biphasic manner with increasing CD44 expression. Simulations using the motor clutch model established that the results could be explained in terms of the strength of cell-substrate interactions. In contrast to these studies, our results show that adhesion strength between cells, mediated by E-cadherin expression, gives rise to the observed non-monotonic behavior in the tumor proliferation (Fig. 2a). The mechanism, identified here, is related to the pressure dependent feedback on growth [15] whose effectiveness is controlled by cell-cell adhesion strength, *f*^*ad*^.

### Comparison With Experiments

We consider three experiments, which provide support to our conclusions: (i) Based on the observation of enhanced cell migration due to lower E-cadherin levels, it has been proposed that tumor invasion and metastasis follows the loss of E-cadherin expression [24]. However, most breast cancer primary and metastatic tumors express E-cadherin [54]. To resolve this discrepancy, Padmanaban et. al [38] compared the tumor growth behavior between E-cadherin negative cells (characterized by reduced E-cadherin expression compared to control) and control E-cadherin expressing cells using three-dimensional (3D) tumor organoids. They report that tumors arising from low E-cadherin expressing cells (E-cad(-)) were smaller than tumors from control E-cadherin (E-cad(control)) expressing cells at corresponding time points over multiple weeks of tumor growth (see Fig. 5a Lower Inset). Using the experimental data from Ref. [38], we extracted the tumor growth rate for E-cad(-) and E-cad(control) organoids, and observe enhanced growth rate for tumors made up of E-cad(control) cells. The longest tumor dimension for E-cadh(control) tumor organoids expanded 1.3 times faster compared to E-cad(-) tumor organoids (see Bottom Inset, Fig. 5a). From our tumor simulations, we predicted the E-cadh(control) tumor spheroids expand 1.8 times faster compared to E-cad(-) tumor spheroids (see Top Inset, Fig. 5a). This shows good agreement between theory and experimental results. It is worth emphasizing that the simulation results were obtained without any fits to the experiments [38], which appeared while this article was already submitted. Comparison of the growth in tumor diameter between E-cad(-) cells and E-cad(control) (red and blue lines respectively, Fig. 5a Main Panel) over 7.5 days of simulated tumor growth shows similar growth rates until *t* ∼ 4 days followed by enhanced growth rates for E-cad(control) cells at *t* > 4 days. The fit for the simulated tumor diameter growth rate is obtained by analyzing data at *t* > 4 days (see Top Inset, Fig. 5a). The tumor size growth rate over multiple days is best fit by a linear function, in agreement between simulation and experiment. The overall magnitude of the tumor growth rate is higher in the experiments as compared to the simulations. The difference in the magnitude of tumor growth rate could be due to the variation in the sizes of cells. In simulations, the maximum cell diameter is 10 *μm* while cells can be larger than 20 *μm* in the experimental tumor spheroids. Other factors such as the difference in cell cycle times, critical pressure which limits growth and spatially asymmetric growth of the tumor could also lead to differences in the overall magnitude of the tumor growth rate. However, both the linear functional form of the tumor growth rates and the higher growth rates in E-cad(control) tumor spheroids as compared to E-cad(-) tumor spheroids are in agreement with simulation predictions.

To conclusively show that the fraction of proliferating cells determine the non-monotonic tumor growth behavior between low E-cadherin expressing tumors and cells expressing control levels of E-cadherin, we turn to further analysis of experimental data from Ref. [38]. By staining tumor cell colonies for PH3 (phospho-histone 3; a mitotic marker), Ref. [38] quantifies the mitotic fraction i.e. the ratio of the number of actively dividing cells to the total number of cells. In agreement with predictions from our simulations, Padmanaban et. al [38] report that E-cadherin expressing tumor cell colonies are indeed characterized by a larger mitotic fraction (blue histogram, Fig. 5b) as opposed to low E-cadherin expressing tumor cells (red histogram, Fig. 5b). We compare the mean mitotic fraction, *F*_*C*_, in E-cad(-) and E-cad(control) in the insets of Fig. 5b between theory and experiment. In agreement with our simulation predictions, enhanced mitotic fraction is seen in tumor spheroids with normal E-cad expression (see Insets of Fig. 5b).

(ii) Maintenance of appropriate E-cadherin expression is required for normal cell development in several species such as drosophila, zebrafish and mouse [44, 55]. We consider a study on how the proliferation capacity of primordial germ cells (PGCs) depend on cell-cell adhesion. Even though the total number of PGCs in an embryo is only around 10-30 cells, while there are thousands of cells in the simulations, we compare the percentage change in cell numbers that result from modulating E-cadherin expression levels. We make this comparison to propose that perhaps E-cadherin does modulate proliferation through a pressure dependent mechanistic pathway in tissues, as discussed above. The proliferation behavior of PGCs in *Xenopus laevis* with changing E-cadherin expression levels is shown in Fig. 5c. In this experiment, E-cadherin overexpression, achieved by mRNA injection of GFP E-cadherin DELE mRNA (E-cad GFP), leads to ∼ 50% reduction (compared to control) in the number of cells after 1.5 days of post fertilization embryo growth [44]. Similar reduction in the number of cells is observed with E-cadherin knockdown using specific morpholino oligonucleotides (E-cad MO) [44]. Therefore, an optimal level of cell-cell adhesion exists where proliferation is maximized, in agreement with simulation predictions (Fig. 5d). The section III in SI provides other examples for the role of E-cadherin in both tumor suppression and proliferation. (iii) The loss of E-cadherin is considered to be a key characteristic in epithelial to mesenchymal transitions, priming the cells to dissociate from the primary tumor and invade surrounding tissues [56]. However, in a subset of high grade glioblastoma, patients with tumor cells expressing E-cadherin correlated with worse prognosis compared to patients whose tumor cells that did not express E-cadherin [36]. In this tumor type, heightened expression of E-cadherin correlated with increased invasiveness in xenograft models [36]. These experimental results, which are consistent with our simulations in promoting proliferation as *f*^*ad*^ is changed from = 0 *μ*N*/μ*m^2^ to 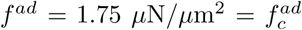 (Fig. 2a), suggest an unexpected role of E-cadherin in promoting tumor growth and invasion.

### Cautionary Remarks

As detailed in the SI section III, it is difficult to make precise comparisons between the simulations and experiments because the growth of tumors is extremely complicated. For example, other cytoplasmic and nuclear signaling factors may be important in understanding the role of cell-cell adhesion in modulating the proliferative capacity of cells, [37] as multiple signaling pathways are located in direct proximity to the adherens junction complexes [20]. Nevertheless, our results suggest that the mechanism of contact inhibition of proliferation, based on critical cellular pressure, could serve as a unifying mechanism in understanding how cell-cell adhesion influences proliferation. The relation between proliferation and cell-cell adhesion has important clinical applications, such as in the development of innovative therapeutic approaches to cancer [57, 58] by targeting E-cadherin expression. Under certain circumstances, inhibiting E-cadherin expression could lead to tumor progression. On the other hand, in cancer cells nominally associated with low or negligible E-Cadherin levels, tumor progression and worsening prognosis could result upon increasing E-Cadherin levels (see SI, Section III for further discussion).

## CONCLUSIONS

In this study, we have established that the modulation of cell-cell adhesion strength contributes to contact inhibition of cell growth and proliferation. Surprisingly, cell proliferation exhibits a non-monotonic behavior as a function of cell adhesion strength, increasing till a critical value, followed by proliferation suppression at higher values of *f*^*ad*^. We have shown that E-cadherin expression and critical pressure based contact inhibition are sufficient to explain the role of cell-cell adhesion on cell proliferation in the context of both morphogenesis and cancer progression. The observed dual role that E-cadherin plays in tumor growth is related to a feedback mechanism due to changes in pressure as the cell-cell interaction strength is varied, established here on the basis of simulations and a mean field theory.

The pressure feedback on the growth of cells is sufficient to account for cell proliferation in the simulations. For cells, however, it may well be that mechanical forces do not directly translate into proliferative effects. Rather, cell-cell contact (experimentally measurable through the contact length *l*_*c*_, for example) could biochemically regulate Rac1/RhoA signaling, which in turn controls proliferation, as observed in biphasic proliferation of cell collectives in both two and three dimensions [39, 40].

One implication of our finding is that the mechanical pressure dependent feedback may also play a role in organ size control. As tissue size regulation requires fine tuning of proliferation rate, cell volume, and cell death at the single cell level [59], pressure dependent feedback mediated by cell-cell adhesion could function as an efficient control parameter. In principle, cells in tissues could be characterized by a range of adhesion strengths. Competition between these cell types, mediated by adhesion dependent pressure feedback into growth, could be critical in determining the relative proportion of cells and therefore the organ size.

## Appendix A: Methods

### Model

We simulate the collective movement of cells using a minimal model of an evolving tumor embedded in a matrix using an agent-based three dimensional (3D) model [60–62]. The cells, which are embedded in a highly viscous material mimicking the extracellular material, are represented as deformable objects. The inter-cell interactions are characterized by direct elastic (repulsive) and adhesive (attractive) forces. The total force on the *i*^*th*^ cell is given by,

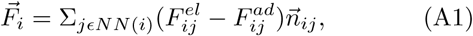

where 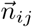 is the unit vector from the center of cell *j* to cell *i*. The forces are summed over the nearest neighbors (*NN* (*i*)) of the *i*^*th*^ cell. The form of the elastic force, 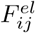 (see Eq. A2), and the inter-cell adhesive force, 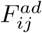 (see Eq. A3), are taken from the study of Schaller and Meyer-Hermann [60]. The strength of the adhesive interaction between cells, *f*^*ad*^, is measured in units of *μN/μm*^2^, (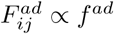 given by Eq. A3). Force as a function of cell center-to-center distance is plotted in Fig. 6c.

Cell-to-cell and cell-to-matrix damping account for the effects of friction due to other cells, and the extracellular matrix (ECM) (for example, collagen matrix), respectively. The model accounts for apoptosis, cell growth and division. Thus, the collective motion of cells is determined by both systematic cell-cell forces and the dynamics due to stochastic cell birth and apoptosis under a free boundary condition [63].

### Forces Between Cells and Equations of Motion

Each cell is represented as a soft sphere whose radius changes in time to account for cell growth. We characterize each cell by its radius, elastic modulus, membrane E-cadherin receptor, and ligand concentration. Following previous studies [60, 61, 64], we used Hertzian contact mechanics to model the magnitude of the elastic force between two spheres of radii *R*_*i*_ and *R*_*j*_, given by,

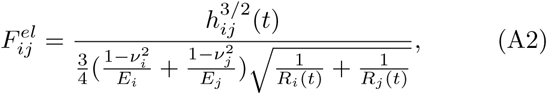

where the parameters *E*_*i*_ and *ν*_*i*_, respectively, are the elastic modulus and Poisson ratio of the *i*^*th*^ cell [60]. The overlap between cells, if they interpenetrate without deformation, is *h*_*ij*_, defined as 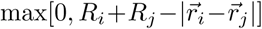 with 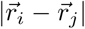 being the center-to-center distance (see Fig. 1b in the Main Text). The elastic repulsive forces tend to minimize the overlap between cells, and could be thought of as a proxy for cortical tension [65, 66].

The magnitude of the attractive adhesive force, 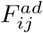, between cells *i* and *j* is given by,

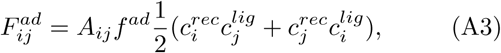

where *A*_*ij*_ is the cell-cell contact area, 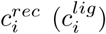 is the E-cadherin receptor (ligand) concentration (assumed to be normalized with respect to the maximum receptor or ligand concentration such that 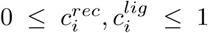). The coupling constant *f*^*ad*^ in Eq. A3, with dimensions *μN/μm*^2^, allows us to rescale the adhesion force, to account for the variations in the maximum receptor and ligand concentrations. Higher (lower) maximum receptor and ligand concentration on the cell surface membrane, is accounted for by higher (lower) value of *f*^*ad*^. It should be noted that the strength of adhesion between the cells is mediated by both the extracellular portion of E-cadherin and how it interacts with the cytoskeleton. The cytoplasmic E-cadherin domain, in conjunction with *α*-catenin, binds to *β*-catenin, linking it to the actin cytoskeleton [66]. In the minimal model, all of these complicated processes that occur on sub-cellular length scales are subsumed in *f*^*ad*^. The inter cell contact surface area, *A*_*ij*_ (see Eq. A3), is obtained using the Hertz model prediction, *A*_*ij*_ = *πh*_*ij*_*R*_*i*_*R*_*j*_*/*(*R*_*i*_ + *R*_*j*_) [60].

An optimal range of cell-cell packing (*h*_*ij*_) exists where *p*_*i*_ is minimized (see Fig. 7b). With the definition *p*_*i*_, growth in the total number of cells (*N* (*t*)) is well approximated as an exponential *N* (*t*) ∝ e*xp*(*const* × *t*) at short times (*t* < 10^5^ secs) (see Fig. 7a). At longer time scales (*t* > ∼ 3 × 10^5^ secs), the increase in the tumor size follows a power law, *N* (*t*) ∝ *t*^*β*^. Such a cross over from exponential to power-law growth in 3D tumor spheroid size has been observed in many tumor cell lines with *β* varying from one to three [67–71]. Based on the power law growth behavior, we expect, 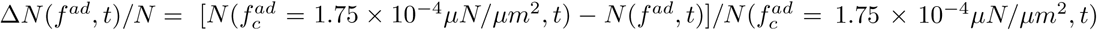, for *f*^*ad*^ = 0 to be 1 (3 × 10^−6^*t*^1.6^)*/*(2.1 × 10^−10^*t*^2.4^). Hence, Δ*N* (*f*^*ad*^ = 0, *t*)*/N* = 1 − *A*_0_*t*^−0.8^, where *A*_0_ = 1.4 × 10^−4^ is a constant. Similarly for Δ*N* (*f*^*ad*^ = 3 × 10^−4^*μN/μm*^2^, *t*)*/N* = 1 − *A*_1_*t*^−1^ where *A*_1_ = 2.2 × 10^5^. Due to the local cell density fluctuations caused by the birth-death events, the cell pressure *p*_*i*_ is a highly dynamic quantity (see Figs. 8a-c). *p*_*i*_(*t*) plays an important role in how local forces provide a feedback on cell growth and division.

While cells are characterized by many different types of cell adhesion molecules (CAMs), here we focus on E-cadherin. We consider different levels of CAM expression, varying from low (*f*^*ad*^ = 0 *μ*N*/μ*m^2^) to intermediate (*f*^*ad*^ = 1.5 *μ*N*/μ*m^2^) to high (*f*^*ad*^ = 3 *μ*N*/μ*m^2^) values. For a discussion on the appropriateness of the value of the range of *f*^*ad*^, see Appendix B. We used a distance sorting algorithm to efficiently obtain a list of nearest neighbors in contact with the *i*^*th*^ cell. For a cell, *i*, an array with distances from cell *i* to all the other cells is created. We then calculated 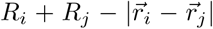 and sorted for cells *j* that satisfy the condition 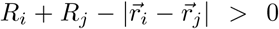, a necessary condition for any cell *j* to be in contact with cell *i*.

The justification that the inertial forces can be neglected can be found in our previous study (see Fig. 18 in Ref. [63]). If we neglect inertial effects, the equation of motion of the *i*^*th*^ cell is,

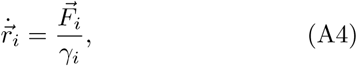

where, 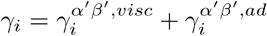 is the friction coefficient with

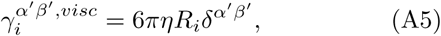

and

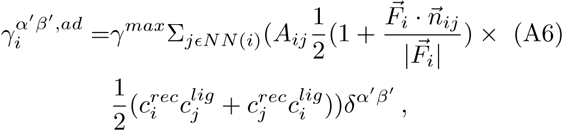

being the cell-to-matrix and cell-to-cell damping contributions respectively. Here, the indices *α*′, *β*′ represent cartesian co-ordinates. Viscosity of the medium surrounding the cell is denoted by *η* and *γ*^*max*^ is the adhesive friction coefficient. Additional details of the simulation methods are given elsewhere [63]. Note that because the equations of motion for the coarse-grained model contain the friction term they do not satisfy Galilean invariance.

### Pressure-dependent Dormancy

A crucial feature in the model is the role played by the local pressure, *p*_*i*_, experienced by the *i*^*th*^ cell relative to a critical pressure, *p*_*c*_. A given cell, at any time *t*, can either be in the dormant (*D*) or in the growth (*G*) phase depending on the pressure on the cell (see Fig 1b). The total pressure (*p*_*i*_) on the *i*^*th*^ cell,

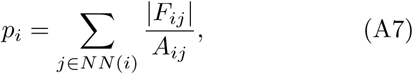

is the overall sum of all the normal pressures due to the nearest neighbors. If *p*_*i*_ > *p*_*c*_ (a pre-assigned value of the critical pressure), the cell becomes dormant, and can no longer grow or divide. Note that a cell that becomes dormant at time *t* does not imply that it remains so at all later times because as the cell colony evolves, *p*_*i*_, a dynamic quantity fluctuates, and hence can become less or greater than *p*_*c*_ (see Figs. 8a-c, Supplementary Movies 1-3A). It has been shown *in vitro* that solid stress, defined as the mechanical stress due to solid and elastic elements of the extracellular matrix, inhibits growth of multicellular tumor spheroids irrespective of the host species, tissue origin or differentiation state [72]. This type of growth inhibition is mediated by stress accumulation around the spheroid as a result of the progressive displacement of the surrounding matrix due to the growing clump of cells. In our model, however, the effect of pressure on cell growth is driven by local cell-cell contact as opposed to the global stress exerted by the surrounding matrix. Both *in vivo* and *in vitro*, epithelial cells exhibit contact inhibition of proliferation due to cell-cell interactions [73]. The value of the critical pressure used in our work is in the same range as experimentally measured cell-scale stresses (10-200 Pa) [49] and with earlier works using critical cellular compression as the mechanism for contact inhibition [60, 62]. We have also verified that the qualitative results are independent of the precise value of *p*_*c*_, as well as alternative definitions of local pressure (see Sections I-II in the SI).

### Cell Dynamics

Because the Reynolds number for cells in a tissue is small [60, 74], overdamped approximation is appropriate. The equations of motion are given below (see Eq. A4). Besides the cell-cell repulsive and adhesive forces, another contribution to cell dynamics comes from cell growth, division and apoptosis. Stochastic cell growth leads to dynamic variations in the cell-cell forces. Cell division and apoptosis induce temporal rearrangements in the cell positions and packing. Hence, both the contribution of the systematic forces and cell growth, birth and apoptosis towards cell dynamics are taken into account in the model, for which we described the unusual dynamics previously [63, 75]. The parameters used in the simulations are given in the SI (see Table I). We justify the range of *f*^*ad*^ explored in our simulations in Appendix B. In order to explore the plausible dual role of E-cadherin on cell proliferation, we vary *f*^*ad*^, keeping all other parameters constant.

## Appendix B: Calibration of Cell-Cell Adhesion Strength

The crucial parameter in the present study is the cell-cell interaction strength, *f*^*ad*^, which is a proxy for E-cadherin expression. In order to assess if the values used in our simulations are in a reasonable range, we estimated *f*^*ad*^ from the typical strength of cell-cell attractive interactions reported in previous studies. Early experiments showed that the interaction strength between cell adhesion proteoglycans is ∼ 2 × 10^−5^*μN/μm*^2^ [76]. More recently, single cell force spectroscopy (SCFS) technique has been used to measure directly the typical forces required to rupture E-cadherin mediated bonds between cells. Several types of cadherins could be present on the cell surface, in addition to adhesion molecules such as integrins, selectins etc [46]. In order to confirm that it is indeed E-Cadherin expression level that changes at different stages of embryo development, Baronsky et. al [44] functionalized gold coated substrate with E-cadherin, and measured the force-distance curves between primordial germ cells and the substrate.

Within the range of *f*^*ad*^ considered in the simulations, a typical force distance curve (plot of 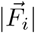 versus 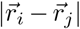 from Eq. A1) is shown in Fig. 6c. The plot in Fig. 6c shows that for typical cell sizes (≈ 5*μm*) the minimum force is ≈ 2 ×10^−4^*μN*, which is fairly close to the values ≈ 1.5 ×10^−4^*μN* in Fig. 6d and 4 × 10^−4^*μN* reported elsewhere [76]. The inset in Fig. 6d shows the work done to overcome the adhesion force mediated by E-cadherin which is the area under the force-distance curve (FDC). The work expended is comparable at 0.78 × 10^−16^Nm for primordial germ cells (PGCs in *Xenopus laevis* embryo) to E-cadherin substrate separation experiment and 1.25 × 10^−16^Nm for cell-cell separation in the model. Because the set up in single cell force spectroscopy and the theoretical model are not precisely comparable, it is gratifying that the magnitude of forces required to separate two cells obtained using Eq. (A1) and the measured values are not significantly different. Note that we did not adjust any parameters to obtain the reasonable agreement. We undertook this comparison to merely point out that the range of E-cadherin mediated forces used in our simulations reflects the typical cell-cell adhesion strength measured in experiments.

The timescale associated with single receptor-ligand binding is typically 2-10 seconds [62]. There are about ∼ 10 cadherins/*μm*^2^ on the surface of typical cells [42], corresponding to ∼ 3000 cadherins on the cell surface for a cell with radius of 5 microns. For studies of cell growth and dynamics on the time scale of days, the fluctuations at the level of single receptor-ligand binding can therefore be neglected [62], thus justifying the use of constant *f*^*ad*^ values. The receptor/ligand concentration are sampled from a Gaussian distribution (see SI Section IV for more details).

Given the center-to-center distance, *r*_*ij*_ = |*r*_*i*_ − *r*_*j*_ |, between cells *i* and *j*, the contact length, *l*_*c*_, and contact angle *β* can be calculated. Let *x* be the distance from center of cell *i* to contact zone marked by *l*_*c*_, along *r*_*ij*_. Similarly, we define *y* as the distance between center of cell *j* to *l*_*c*_ once again along *r*_*ij*_ (see Fig. 1a of Main Text). Based on the right triangle that is formed between *x, R*_*i*_ and 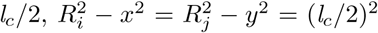 and *x* + *y* = *r*_*ij*_. This allows us to solve for *x, y* and hence,

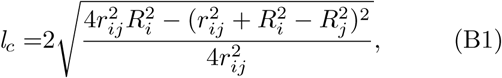

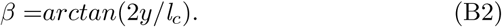

The probability distribution for *l*_*c*_ and *β* obtained from the simulation for varying values of *f*^*ad*^ is shown in Figs. 6a - 6b.

The interaction parameters characterizing the model are, the two elastic constants (*E*_*i*_ and *ν*_*i*_) and *f*^*ad*^ if we assume that the combination of receptor and ligand concentrations in Eq. (A3) is a constant. In addition, the evolution of cell colony introduces two other parameters, birth (*k*_*b*_) and apoptotic rates (*k*_*a*_) of cells. The values of *k*_*a*_ and *k*_*b*_ depend on the detailed biology governing cell fate, which we simply take as parameters in the simulations. If the elastic constants, and *k*_*a*_, *k*_*b*_ are fixed, then the only parameter that determines the evolution of the tumor is *f*^*ad*^ and *p*_*c*_, whose magnitude is determined by E-cadherin expression. Here, we explore the effects of *f*^*ad*^ and *p*_*c*_, on tumor proliferation.

## Appendix C: Average Number of Nearest Neighbor of Cells Increases with *f*^*ad*^

The collective movement of cells (related to proliferative capacity) is determined by cell arrangement and packing within the three dimensional (3D) tissue, which clearly depends on the adhesion strength. Analyzing the distribution of number of nearest neighbors (see Fig. 9a), the arrangement of cells in the spheroid has a peak near 6 nearest neighbors at low *f*^*ad*^. Very few cells, if any, have less than 2 neighbors. Similarly, at *f*^*ad*^ = 5 × 10^−5^ *μ*N*/μ*m^2^, few cells have more than 9 neighbors (see Fig. 9c). As *f*^*ad*^ increases, the peak in the nearest neighbor distribution moves to higher values, with the distribution also becoming broader (Fig. 9a). With the highest adhesion strength, *f*^*ad*^ = 3 × 10^−4^*μ*N*/μ*m^2^, the average number of nearest neighbors is ≈ 9 cells which is consistent with 3D experimental data for mouse blastocyst after 5 − 9 days of growth [78].

We surmise that the dependence of the average number of nearest neighbors on cell-cell adhesion strengths *f*^*ad*^ in the simulations is consistent with experimental findings. Cell packing data in 2D epithelial structures, quantified by the probability distribution of nearest neighbors [77], allow us to compare the simulation results to experiments (Fig. 9c). We compare the simulation results for the distribution of the number of nearest neighbors (*n*_*NN*_) with experiment, keeping in mind that our simulation is in 3D. In 3D, the average number of nearest neighbors is higher than in 2D. There is indeed an increase in the probability of nearest neighbors from 7 − 10 (Fig. 9c). Moreover, *f*^*ad*^ > 100 dynes*/*cm^2^ = 10^−5^*μ*N*/μ*m^2^, within an order of magnitude has been reported in experiments [79], which we point out only to show that the values of *f*^*ad*^ considered in our study are reasonable.

The average number of nearest neighbors, 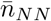, increases as *f*^*ad*^ increases (Fig. 9b). The approximate linear fit 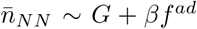 is used to rationalize the data in Fig. 2a in the Main Text. The fit parameters *G, β* are given in the caption. The linear fit is used only for calculating 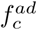, the optimal value at which proliferation is a maximum.

### Distribution of h_*ij*_

The cell-cell overlap, *h*_*ij*_, gives an indication of how closely the deformable cells are packed within the 3D spheroid. At low adhesion strengths, the distribution is sharply peaked at small *h*_*ij*_ (see Fig. 10a), implying there is minimal cell-cell interpenetration. As *f*^*ad*^ increases, the cells are jammed. For *f*^*ad*^ = 3 × 10^−4^ *μ*N*/μ*m^2^, the average cell overlap 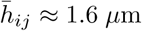, implying that the center to center distance between cells is approximately 6.4 *μ*m (for cells of radii 4 *μ*m). Note that the cell overlap distribution becomes broader as *f*^*ad*^ increases. The mean overlap, 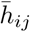, varies quadratically with adhesion strength (Fig. 10b). If we set *F* ^*el*^ = *F*^*ad*^, we find that *h* ∼ (*f*^*ad*^)^2^, and as expected we obtain the fit 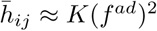.

To calculate the total pressure experienced by a cell (*p*_*t*_) theoretically, as detailed in the Main text, we look at the deviation of the average cell overlap 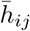 from the optimal overlap (*h*_0_) as *f*^*ad*^ is changed, where *h*_0_ is the overlap distance at which the pressure experienced by a cell is a minimum. This would occur when the repulsive and attractive interaction forces between a pair of cells balance (*F*^*el*^ = *F*^*ad*^). The deviation, 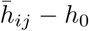, increases as *f*^*ad*^ increases, an indication that it is harder for cells to relax to optimal intercellular distances, due to packing frustration. For the purposes of rationalizing the optimal value of *f*^*ad*^ (see Main Text) we write, 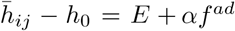, where the parameters *E, α* are as listed in Fig. 11. We found that *h*_0_ depends on radii of the cells that are in contact as well as other parameters,

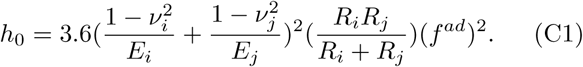

Hence, we estimate 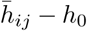 by using average quantities.

## Acknowledgements

We acknowledge Anne D. Bowen at the Visualization Laboratory (Vislab), Texas Advanced Computing Center, for help with video visualizations. This work is supported by the National Science Foundation (PHY 17-08128 and CHE 16-32756) and the Collie-Welch Chair through the Welch Foundation (F-0019).

## References

[1] Loïc LeGoff and Thomas Lecuit. Mechanical forces and growth in animal tissues. Cold Spring Harbor perspectives in biology, 8(3):a019232, 2016.

[2] Peter Friedl and Roberto Mayor. Tuning collective cell migration by cell–cell junction regulation. Cold Spring Harbor perspectives in biology, 9(4):a029199, 2017.

[3] Srikanth Budnar and Alpha S Yap. A mechanobiological perspective on cadherins and the actin-myosin cytoskeleton. F1000prime reports, 5(35):1–6, 2013.

[4] Dhananjay T Tambe, C Corey Hardin, Thomas E Angelini, Kavitha Rajendran, Chan Young Park, Xavier Serra-Picamal, Enhua H Zhou, Muhammad H Zaman, James P Butler, David A Weitz, et al. Collective cell guidance by cooperative intercellular forces. Nature materials, 10(6):469, 2011.

[5] Jennifer M Halbleib and W James Nelson. Cadherins in development: cell adhesion, sorting, and tissue morphogenesis. Genes & development, 20(23):3199–3214, 2006.

[6] Masatoshi Takeichi. The cadherins: cell-cell adhesion molecules controlling animal morphogenesis. Development, 102(4):639–655, 1988.

[7] Masataka Saito, Dana K Tucker, Drew Kohlhorst, Carien M Niessen, and Andrew P Kowalczyk. Classical and desmosomal cadherins at a glance. J Cell Sci, 125(11):2547–2552, 2012.

[8] Erdem Tabdanov, Nicolas Borghi, Françoise Brochard-Wyart, Sylvie Dufour, and Jean-Paul Thiery. Role of e-cadherin in membrane-cortex interaction probed by nanotube extrusion. Biophysical journal, 96(6):2457–2465, 2009.

[9] Nicolas Borghi, Maria Sorokina, Olga G Shcherbakova, William I Weis, Beth L Pruitt, W James Nelson, and Alexander R Dunn. E-cadherin is under constitutive actomyosin-generated tension that is increased at cell–cell contacts upon externally applied stretch. Proceedings of the National Academy of Sciences, 109(31):12568–12573, 2012.

[10] Darcy Wentworth Thompson et al. On growth and form. Cambridge Univ. Press, 1942.

[11] Tanya J Shaw and Paul Martin. Wound repair at a glance. Journal of cell science, 122(18):3209–3213, 2009.

[12] M Abercrombie. Contact inhibition in tissue culture. In vitro, 6(2):128–142, 1970.

[13] Judah Folkman and Anne Moscona. Role of cell shape in growth control. Nature, 273(5661):345, 1978.

[14] Christopher S Chen, Milan Mrksich, Sui Huang, George M Whitesides, and Donald E Ingber. Geometric control of cell life and death. Science, 276(5317):1425–1428, 1997.

[15] Boris I Shraiman. Mechanical feedback as a possible regulator of tissue growth. Proceedings of the National Academy of Sciences of the United States of America, 102(9):3318–3323, 2005.

[16] Sebastian J Streichan, Christian R Hoerner, Tatjana Schneidt, Daniela Holzer, and Lars Hufnagel. Spatial constraints control cell proliferation in tissues. Proceedings of the National Academy of Sciences, 111(15):5586–5591, 2014.

[17] Shane Jacobeen, Jennifer T Pentz, Elyes C Graba, Colin G Brandys, William C Ratcliff, and Peter J Yunker. Cellular packing, mechanical stress and the evolution of multicellularity. Nature Physics, 14(3):286, 2018.

[18] Alberto Puliafito, Lars Hufnagel, Pierre Neveu, Sebastian Streichan, Alex Sigal, D Kuchnir Fygenson, and Boris I Shraiman. Collective and single cell behavior in epithelial contact inhibition. Proceedings of the National Academy of Sciences, 109(3):739–744, 2012.

[19] Andrea I McClatchey and Alpha S Yap. Contact inhibition (of proliferation) redux. Current opinion in cell biology, 24(5):685–694, 2012.

[20] Antonis Kourtidis, Ruifeng Lu, Lindy J Pence, and Panos Z Anastasiadis. A central role for cadherin signaling in cancer. Experimental cell research, 358(1):78–85, 2017.

[21] Kenneth D Irvine and Boris I Shraiman. Mechanical control of growth: ideas, facts and challenges. Development, 144(23):4238–4248, 2017.

[22] Kris Vleminckx, Luc Vakaet, Marcus Mareel, Walter Fiers, and Frans Van Roy. Genetic manipulation of E-cadherin expression by epithelial tumor cells reveals an invasion suppressor role. Cell, 66(1):107–119, 1991.

[23] Anne-Karina Perl, Petra Wilgenbus, Ulf Dahl, Henrik Semb, and Gerhard Christofori. A causal role for E-cadherin in the transition from adenoma to carcinoma. Nature, 392(6672):190, 1998.

[24] Uwe H Frixen, Jfirgen Behrens, Martin Sachs, Gertrud Eberle, Beate Voss, Angelika Warda, Dorothea Löchner, and Walter Birchmeier. E-cadherin-mediated cell-cell adhesion prevents invasiveness of human carcinoma cells. The Journal of cell biology, 113(1):173–185, 1991.

[25] Alice ST Wong and Barry M Gumbiner. Adhesion-independent mechanism for suppression of tumor cell invasion by E-cadherin. The Journal of cell biology, 161(6):1191–1203, 2003.

[26] Barry M Gumbiner. Cell adhesion: the molecular basis of tissue architecture and morphogenesis. Cell, 84(3):345–357, 1996.

[27] Amparo Cano, Mirna A Pérez-Moreno, Isabel Rodrigo, Annamaria Locascio, María J Blanco, Marta G del Barrio, Francisco Portillo, and M Angela Nieto. The transcription factor snail controls epithelial-mesenchymal transitions by repressing E-cadherin expression. Nature cell biology, 2(2):76, 2000.

[28] Jing Yang and Robert A Weinberg. Epithelial-mesenchymal transition: at the crossroads of development and tumor metastasis. Developmental cell, 14(6):818–829, 2008.

[29] Robert Weinberg. The biology of cancer. Garland science, New York, 2013.

[30] Yuanmei Lou, Olena Preobrazhenska, Margaret Sutcliffe, Lorena Barclay, Paul C McDonald, Calvin Roskelley, Christopher M Overall, Shoukat Dedhar, et al. Epithelial-mesenchymal transition (EMT) is not sufficient for spontaneous murine breast cancer metastasis. Developmental Dynamics, 237(10):2755–2768, 2008.

[31] Kari R Fischer, Anna Durrans, Sharrell Lee, Jianting Sheng, Fuhai Li, Stephen TC Wong, Hyejin Choi, Tina El Rayes, Seongho Ryu, Juliane Troeger, et al. Epithelial-to-mesenchymal transition is not required for lung metastasis but contributes to chemoresistance. Nature, 527(7579):472, 2015.

[32] Xiaofeng Zheng, Julienne L Carstens, Jiha Kim, Matthew Scheible, Judith Kaye, Hikaru Sugimoto, Chia-Chin Wu, Valerie S LeBleu, and Raghu Kalluri. Epithelial-to-mesenchymal transition is dispensable for metastasis but induces chemoresistance in pancreatic cancer. Nature, 527(7579):525, 2015.

[33] Kevin J Cheung, Edward Gabrielson, Zena Werb, and Andrew J Ewald. Collective invasion in breast cancer requires a conserved basal epithelial program. Cell, 155(7):1639–1651, 2013.

[34] Eliah R Shamir, Elisa Pappalardo, Danielle M Jorgens, Kester Coutinho, Wen-Ting Tsai, Khaled Aziz, Manfred Auer, Phuoc T Tran, Joel S Bader, and Andrew J Ewald. Twist1-induced dissemination preserves epithelial identity and requires E-cadherin. J Cell Biol, 204(5):839–856, 2014.

[35] Deborah Silvera, Rezina Arju, Farbod Darvishian, Paul H Levine, Ladan Zolfaghari, Judith Goldberg, Tsivia Hochman, Silvia C Formenti, and Robert J Schneider. Essential role for eIF4GI overexpression in the pathogenesis of inflammatory breast cancer. Nature cell biology, 11(7):903, 2009.

[36] Laura J Lewis-Tuffin, Fausto Rodriguez, Caterina Giannini, Bernd Scheithauer, Brian M Necela, Jann N Sarkaria, and Panos Z Anastasiadis. Misregulated E-cadherin expression associated with an aggressive brain tumor phenotype. PloS one, 5(10):e13665, 2010.

[37] Fausto J Rodriguez, Laura J Lewis-Tuffin, and Panos Z Anastasiadis. E-cadherin’s dark side: possible role in tumor progression. Biochimica et Biophysica Acta (BBA)-Reviews on Cancer, 1826(1):23–31, 2012.

[38] Veena Padmanaban, Ilona Krol, Yasir Suhail, Barbara M Szczerba, Nicola Aceto, Joel S Bader, and Andrew J Ewald. E-cadherin is required for metastasis in multiple models of breast cancer. Nature, 573(7774):439–444, 2019.

[39] Wendy F Liu, Celeste M Nelson, Dana M Pirone, and Christopher S Chen. E-cadherin engagement stimulates proliferation via Rac1. The Journal of cell biology, 173(3):431–441, 2006.

[40] Darren S Gray, Wendy F Liu, Colette J Shen, Kiran Bhadriraju, Celeste M Nelson, and Christopher S Chen. Engineering amount of cell–cell contact demonstrates biphasic proliferative regulation through RhoA and the actin cytoskeleton. Experimental cell research, 314(15):2846–2854, 2008.

[41] Alpha S Yap, Guillermo A Gomez, and Robert G Parton. Adherens junctions revisualized: Organizing cadherins as nanoassemblies. Developmental cell, 35(1):12–20, 2015.

[42] Jeffrey D Amack and M Lisa Manning. Knowing the boundaries: extending the differential adhesion hypothesis in embryonic cell sorting. Science, 338(6104):212–215, 2012.

[43] Robert David, Olivia Luu, Erich W Damm, Jason WH Wen, Martina Nagel, and Rudolf Winklbauer. Tissue cohesion and the mechanics of cell rearrangement. Development, 141(19):3672–3682, 2014.

[44] Thilo Baronsky, Aliaksandr Dzementsei, Marieelen Oelkers, Juliane Melchert, Tomas Pieler, and Andreas Janshoff. Reduction in E-cadherin expression fosters migration of Xenopus laevis primordial germ cells. Integrative Biology, 8(3):349–358, 2016.

[45] Angela M Jimenez Valencia, Pei-Hsun Wu, Osman N Yogurtcu, Pranay Rao, Josh DiGiacomo, Inäs Godet, Lijuan He, Meng-Horng Lee, Daniele Gilkes, Sean X Sun, et al. Collective cancer cell invasion induced by coordinated contractile stresses. Oncotarget, 6(41):43438, 2015.

[46] Frans Van Roy and Geert Berx. The cell-cell adhesion molecule E-cadherin. Cellular and molecular life sciences, 65(23):3756–3788, 2008.

[47] Andre Glebovich Kamkin and Irina S Kiseleva. Mechanosensitivity in cells and tissues. Springer, 2005.

[48] ME Dolega, Morgan Delarue, François Ingremeau, Jacques Prost, Antoine Delon, and Giovanni Cappello. Cell-like pressure sensors reveal increase of mechanical stress towards the core of multicellular spheroids under compression. Nature communications, 8:14056, 2017.

[49] Alessandro Mongera, Payam Rowghanian, Hannah J Gustafson, Elijah Shelton, David A Kealhofer, Emmet K Carn, Friedhelm Serwane, Adam A Lucio, James Giammona, and Otger Campàs. A fluid-to-solid jamming transition underlies vertebrate body axis elongation. Nature, 561(7723):401, 2018.

[50] PA DiMilla, Kenneth Barbee, and DA Lauffenburger. Mathematical model for the effects of adhesion and mechanics on cell migration speed. Biophysical journal, 60(1):15–37, 1991.

[51] Paul A DiMilla, Julie A Stone, John A Quinn, Steven M Albelda, and Douglas A Lauffenburger. Maximal migration of human smooth muscle cells on fibronectin and type IV collagen occurs at an intermediate attachment strength. The Journal of cell biology, 122(3):729–737, 1993.

[52] Hossein Ahmadzadeh, Marie R Webster, Reeti Behera, Angela M Jimenez Valencia, Denis Wirtz, Ashani T Weeraratna, and Vivek B Shenoy. Modeling the two-way feedback between contractility and matrix realignment reveals a nonlinear mode of cancer cell invasion. Proceedings of the National Academy of Sciences, 114(9):E1617–E1626, 2017.

[53] Rebecca L Klank, Stacy A Decker Grunke, Benjamin L Bangasser, Colleen L Forster, Matthew A Price, David J Odde, Karen S SantaCruz, Steven S Rosenfeld, Peter Canoll, Eva A Turley, et al. Biphasic dependence of glioma survival and cell migration on CD44 expression level. Cell reports, 18(1):23–31, 2017.

[54] Christopher I Li, Benjamin O Anderson, Janet R Daling, and Roger E Moe. Trends in incidence rates of invasive lobular and ductal breast carcinoma. Jama, 289(11):1421–1424, 2003.

[55] Brian E Richardson and Ruth Lehmann. Mechanisms guiding primordial germ cell migration: strategies from different organisms. Nature reviews Molecular cell biology, 11(1):37, 2010.

[56] Alisha M Mendonsa, Tae-Young Na, and Barry M Gumbiner. E-cadherin in contact inhibition and cancer. Oncogene, 37:4769–4780, 2018.

[57] Sabine M Brouxhon, Stephanos Kyrkanides, Xiaofei Teng, Veena Raja, M Kerry O’Banion, Robert Clarke, Stephen Byers, Andrew Silberfeld, Carmen Tornos, and Li Ma. Monoclonal antibody against the ectodomain of E-cadherin (DECMA-1) suppresses breast carcinogenesis: involvement of the HER/PI3K/Akt/mTOR and IAP pathways. Clinical Cancer Research, 19(12):3234–3246, 2013.

[58] Patrícia Carneiro, Joana Figueiredo, Renata Bordeira-Carriço, Maria Sofia Fernandes, Joana Carvalho, Carla Oliveira, and Raquel Seruca. Therapeutic targets associated to E-cadherin dysfunction in gastric cancer. Expert opinion on therapeutic targets, 17(10):1187–1201, 2013.

[59] Romain Levayer, Carole Dupont, and Eduardo Moreno. Tissue crowding induces caspase-dependent competition for space. Current Biology, 26(5):670–677, 2016.

[60] Gernot Schaller and Michael Meyer-Hermann. Multicellular tumor spheroid in an off-lattice Voronoi-Delaunay cell model. Physical Review E, 71(5):051910, 2005.

[61] Dirk Drasdo and Stefan Hohme. A single-cell-based model of tumor growth in vitro: monolayers and spheroids. Physical biology, 2(3):133, 2005.

[62] Jörg Galle, Markus Loeffler, and Dirk Drasdo. Modeling the effect of deregulated proliferation and apoptosis on the growth dynamics of epithelial cell populations in vitro. Biophysical journal, 88(1):62–75, 2005.

[63] Abdul N Malmi-Kakkada, Xin Li, Himadri S Samanta, Sumit Sinha, and Dave Thirumalai. Cell growth rate dictates the onset of glass to fluidlike transition and long time superdiffusion in an evolving cell colony. Physical Review X, 8(2):021025, 2018.

[64] P Pathmanathan, Jonathan Cooper, Alexander Fletcher, Gary Mirams, P Murray, J Osborne, Joe Pitt-Francis, A Walter, and SJ Chapman. A computational study of discrete mechanical tissue models. Physical biology, 6(3):036001, 2009.

[65] G Wayne Brodland. The differential interfacial tension hypothesis (DITH): a comprehensive theory for the self-rearrangement of embryonic cells and tissues. Journal of biomechanical engineering, 124(2):188–197, 2002.

[66] Takashi Hayashi and Richard W Carthew. Surface mechanics mediate pattern formation in the developing retina. Nature, 431(7009):647–652, 2004.

[67] Alan D Conger and Marvin C Ziskin. Growth of mammalian multicellular tumor spheroids. Cancer Res, 43(2):556–560, 1983.

[68] Emmanuel Mandonnet, Jean-Yves Delattre, Marie-Laure Tanguy, Kristin R Swanson, Antoine F Carpentier, Hugues Duffau, Philippe Cornu, Rémy Van Effenterre, Ellsworth C Alvord, and Laurent Capelle. Continuous growth of mean tumor diameter in a subset of grade II gliomas. Ann Neurol, 53(4):524–528, 2003.

[69] Monica Simeoni, Paolo Magni, Cristiano Cammia, Giuseppe De Nicolao, Valter Croci, Enrico Pesenti, Massimiliano Germani, Italo Poggesi, and Maurizio Rocchetti. Predictive pharmacokinetic-pharmacodynamic modeling of tumor growth kinetics in xenograft models after administration of anticancer agents. Cancer Res, 64(3):1094–1101, 2004.

[70] D Hart, E Shochat, and Z Agur. The growth law of primary breast cancer as inferred from mammography screening trials data. Br J Cancer, 78(3):382–387, 1998.

[71] David Robert Grimes, Pavitra Kannan, Alan McIntyre, Anthony Kavanagh, Abul Siddiky, Simon Wigfield, Adrian Harris, and Mike Partridge. The role of oxygen in avascular tumor growth. PloS one, 11(4):e0153692, 2016.

[72] Gabriel Helmlinger, Paolo A Netti, Hera C Lichtenbeld, Robert J Melder, and Rakesh K Jain. Solid stress inhibits the growth of multicellular tumor spheroids. Nature biotechnology, 15(8):778–783, 1997.

[73] Harry Eagle and Elliot M Levine. Growth regulatory effects of cellular interaction. Nature, 213(5081):1102–1106, 1967.

[74] John C Dallon and Hans G Othmer. How cellular movement determines the collective force generated by the Dictyostelium discoideum slug. Journal of theoretical biology, 231(2):203–222, 2004.

[75] Himadri S Samanta and D Thirumalai. Origin of superdiffusive behavior in a class of nonequilibrium systems. Physical Review E, 99(3):032401, 2019.

[76] U Dammer, O Popescu, P Wagner, D Anselmetti, HJ Guntherodt, and GN Misevic. Binding strength between cell-adhesion proteoglycans measured by atomic-force microscopy. Science, 267(5201):1173–1175, 1995.

[77] Matthew C Gibson, Ankit B Patel, Radhika Nagpal, and Norbert Perrimon. The emergence of geometric order in proliferating metazoan epithelia. Nature, 442(7106):1038, 2006.

[78] Sabine C Fischer, Elena Corujo-Simon, Joaquin Lilao-Garzon, Ernst HK Stelzer, and Silvia Munoz-Descalzo. Three-dimensional cell neighbourhood impacts differentiation in the inner mass cells of the mouse blastocyst. bioRxiv, page 159301, 2017.

[79] Stephen W Byers, Connie L Sommers, Becky Hoxter, Arthur M Mercurio, and Aydin Tozeren. Role of E-cadherin in the response of tumor cell aggregates to lymphatic, venous and arterial flow: measurement of cell-cell adhesion strength. Journal of cell science, 108(5):2053–2064, 1995.

