## Supplementary Information for "Dual Role of Cell-Cell Adhesion In Tumor Suppression and Proliferation Due to Collective Mechanosensing"

(Dated: February 10, 2020)

### I. NONLINEAR PROLIFERATION BEHAVIOR IS ROBUST TO ALTERNATIVE VALUES OF THE CRITICAL PRESSURE

The pressure experienced by the cell, $p_i = \sum_{j \in NN(i)} \frac{|F_{ij}|}{A_{ij}}$, considers the absolute value of the force $|F_{ij}|$ exerted on a cell $i$. The use of the absolute value ensures that both repulsive (positive) and adhesive (negative) contributions to the pressure are treated on an equal footing, given any fixed positive value of the critical pressure.

In order to ascertain if the non-monotonic dependence of proliferation on $f^{ad}$ depends on exact values and definition of the critical pressure, we varied $p_c$. We also considered alternative definitions of the critical pressure because the precise calculation of pressure in systems that are far from equilibrium is not entirely clear.

**Role of $\mathbf{p}_c$:** For $p_c = 5 \times 10^{-5}$MPa, the size of the spheroid, N(t=7.5 days), is shown in Fig. S1. The biphasic behavior also persists at the lower critical pressure of $p_c = 5 \times 10^{-5}$ MPa, indicating that the non-monotonic behavior of the proliferative capacity as a function of $f^{ad}$ does not depend on the exact value of $p_c$ in the range explored here. However, the proliferation extent measured by the total number of cells obtained after 7.5 days ($\sim 12\tau_{min}$) of growth, at all values of $f^{ad}$, is greatly reduced at lower $p_c$ (compare to Fig. 2a in the Main Text). For example, at $f^{ad} = 1.5 \times 10^{-4} \mu\text{N}/\mu\text{m}^2$ (for $p_c = 5 \times 10^{-5}$MPa), there is a $\approx 40\%$ reduction in $N$ as compared to $p_c = 1 \times 10^{-4}$MPa (at fixed $t$). Lower critical pressure makes it easier for cells to enter the dormant state, leading to the inhibition of overall proliferation while preserving the variation of the size of the tumor as a function of $f^{ad}$.

**Irving-Kirkwood Pressure:** Pressure experienced by cells can also be calculated based on the Irving-Kirkwood (IK) stress tensor. In systems out of equilibrium, it is beneficial to calculate local stress at certain points in space and time. The IK stress tensor (1, 2), $\sigma^{\alpha'\beta'}$, is defined as,

$$\sigma_i^{\alpha'\beta'} = \frac{1}{V} \sum_{i \neq j \in NN(i)} F_{ij}^{\alpha'} |r_i^{\beta'} - r_j^{\beta'}|, \tag{S.1}$$

where $\alpha', \beta' = [x, y, z]$, $V = (4/3)\pi R_i^3 + \sum_{j \in NN(i)} (4/3)\pi R_j^3$ is the local volume occupied by a cell and its nearest neighbors, and $F_{ij}$ is the magnitude of the force exerted on cell $i$ due to cell $j$. Here, the nearest neighbors of a cell $i$ ($NN(i)$) is defined as any cell $j$ with $h_{ij} > 0$. The IK pressure is the trace of the stress tensor $p_i^{IK} = \sigma^{\alpha'\alpha'}/3$.

Non-monotonic proliferation behavior is observed even when the IK definition of pressure (see Fig. S2) is used. The peak in the number of cells is at $f^{ad} \sim 1.75 \times 10^{-4}$, which is fairly close to the value found in Fig. 2 of the Main Text. We find that the magnitude of the pressure calculated using Eq. (S.1) is less than the values obtained by calculating $p_i$ as described in the Main Text. As a result, we used lower values of $p_c$ in order to explore the dependence of the proliferation on the $f^{ad}$. Because $p_c$ is small, the number of cells obtained after 5.6 days of growth is less than what is found in Fig. 2a of the Main Text.

### II. USE OF DIFFERENT FORM OF $f^{ad}$ PRESERVES NON-MONOTONIC PROLIFERATION

Besides ensuring that the nonlinear proliferation behavior does not depend on the value of the critical pressure (see Fig. S1), we also tested whether our results are dependent on the form of the cell-cell interaction (see Eqs. A3- A4 Appendix A of the Main Text). We performed additional simulations using adhesive interaction of the form,

$$F^{ad} = \rho_m W_s h_{ij}, \tag{S.2}$$

where $\rho_m$ is the density of surface adhesion molecules, and $W_s$ is the adhesion energy of a single bond (3). According to Eq. S.2, with decreasing cell center-to-center distance or equivalently increasing cell-cell overlap $h_{ij}$, the number of adhesive contacts between cells increase. This leads to increased attractive interaction between the cells. The repulsive interaction is left unchanged. Defining a new cell-cell adhesion strength parameter, $f^{ad}_{new} = \rho_m W_s$, the biphasic cell proliferation behavior is once again obtained (see Fig. S3). Although the optimal cell-cell adhesion strength shifts to a different value, $f^{ad}_{opt,new} = 6 \times 10^{-4}$ $\mu$N/$\mu$m, compared to the interaction considered in the Main text, the overall trend is similar. Thus, the qualitative observation of non-monotonic dependence of proliferative capacity on $f^{ad}$ is unchanged, establishing the robustness of the results.

### III. PLAUSIBLE CONNECTION BETWEEN SIMULATIONS AND CLINICAL DATA

E-cadherin is considered to be primarily a tumor suppressor, based on the observation that it is down regulated during epithelial to mesenchymal transition (EMT) (4, 5). The tumor suppressor role of E-cadherin (encoded by the CDH1 gene) has been elucidated in breast cancer where loss of heterozygosity in chromosome region 16q22.1 (the gene region that codes for E-cadherin) is frequent (4, 5). In recent years, however, an alternative role for E-cadherin as a tumor promoter seems to be emerging (5–7).

According to our findings (albeit using only simulations), the overall E-cadherin expression level determines its role as a tumor suppressor or promotor. For cells characterized by low/no E-cadherin expression, we hypothesize that increasing its expression leads to enhanced tumorigenicity, and by implication poor survival prognosis. On the other hand, if the native E-cadherin expression level is high, its up-regulation could lead to the suppression of tumor growth. In the context of cancer, heterogeneity in E-cadherin expression is observed. For example, E-cadherin expression is rare to non-existent in both brain tumors and normal brain tissues (8–10). In colorectal tissues, however, epithelial cells express E-cadherin without exception (11, 12). We should point out that it is difficult to quantitatively compare the key prediction made here (Fig. 2 in the Main Text) expressing the dual role of E-cadherin in tumor growth with available data. Nevertheless, it appears that there is evidence for the non-monotonic cell proliferation as a function of the strength of cell-cell attraction. We give anecdotal evidence for the dual role that E-cadherin plays in tumor evolution.

**E-cadherin expression correlates with worse prognosis for glioblastoma and ovarian cancer:** E-cadherin levels in tumor tissue samples from 27 individuals with a rare subtype of Glioblastoma Multiforme (GBM) with epithelial/ pseudoepithelial differentiation was analyzed by Lewis-Tuffin et. al. (8). Nine out of the 27 cases exhibited E-cadherin expression. These patients demonstrated poorer overall survival compared to the 18 patients whose tumors did not express E-cadherin (see Fig. S4a, Negative stands for no E-cadherin expression compared to tumor cells exhibiting Membranous/cytoplasmic E-cadherin expression).

After establishing orthotropic xenografts in mice, Lewis-Tuffin et.al (8) sectioned the brains to determine the relative invasiveness of the tumors depending on E-cadherin expression. Five out of the eight high/moderately invasive tumors expressed enhanced levels of E-cadherin while none of the minimally invasive tumors showed E-cadherin expression (see Fig. S4b). The data indicate that higher E-cadherin expression could be one of the contributors to GBM tumor aggressiveness.

Similar to GBM, E-cadherin expression in ovarian cancers exhibits a distinct pattern. Healthy ovarian surface epithelial (OSE) cells do not express E-cadherin. However, it is consistently expressed in benign, borderline and malignant ovarian tumors at all stages, including in metastases from such ovarian tumors (13, 14). To ascertain the physiological function of E-cadherin in ovarian tumor cells, 3-(4,5-dimethylthiazol-2-yl)-2,5-diphenyltetrazolium bromide (MTT) cell viability and proliferation assays were performed. MTT assay involved seeding of $2 \times 10^5$ of ovarian cancer cell line (OVCAR-3) in 24-well plates with or without E-cadherin neutralizing antibody. Number of cells at time 0 was defined as 1.0, and fold reductions/increments at different time points is indicated as mean $\pm$ standard deviation (14). Control group of cells show consistent proliferation while neutralization of E-cadherin function in cancer cells led to marked suppression of cell proliferation (Fig. S4c). Therefore, one could surmise that E-cadherin plays the role of a tumor promoter in certain forms of Ovarian cancers. As mentioned earlier, mapping our findings to the specific cancer types discussed here requires more precise data analysis, which would require additional experiments and simulations. The qualitative similarity between our findings and in certain cancer types is encouraging.

**Tumor suppression:** In colorectal cancers, infiltrative tumor growth and lymph node metastasis are correlated with loss of E-cadherin expression (15), implying that enhanced $f^{ad}$ leads to tumor suppression. Given the heterogeneity associated with cancer cell properties, it is possible that cells within a single cancer type may exhibit a wide variation in cellular adhesion molecule expression levels. In such a scenario, it would be difficult to isolate the effect of cell-cell adhesion on proliferation. The prediction from our model in terms of tumor proliferation behavior is borne out by the limited analysis we have carried out i.e. role of E-cadherin as a tumor promoter or suppressor depends on the level of gene expression.

### IV. SIMULATION DETAILS

At time, $t = 0$, we begin with seeding 100 cells. The radii of these initial cells are distributed according to a Gaussian, $p(R_i) = \frac{1}{0.5\sqrt{2\pi}} e^{-(R_i-4.5)^2/2\times 0.5^2}$. Similarly, the elastic moduli, $E_i$ and the Poisson ratio $\nu_i$ are also characterized by a Gaussian distribution with standard deviation of $10^{-4}$MPa and 0.02 respectively (mean values are given in Table S1). The receptor and ligand concentration on the cell surface are distributed according to a Gaussian $(p(c_i^{rec}(c_i^{lig})) = \frac{1}{0.02\sqrt{2\pi}} e^{-(c_i^{rec}(c_i^{lig})-0.9)^2/2\times 0.02^2})$, centered around the mean (=0.9) with a dispersion of 0.02. For subsequent cell division cycles, the newborn cell properties are sampled from the same distribution as detailed above. At each time step, the growth rate of the cell is also picked from a Gaussian distribution.

The volume of growing cells increases at a constant rate $r_V$. Cell radii are updated from a Gaussian distribution with the mean rate $\dot{R} = (4\pi R^2)^{-1} r_V$ and dispersion of $10^{-5}$. Over the cell cycle time $\tau$,

$$r_V = \frac{2\pi (R_m)^3}{3\tau}, \tag{S.3}$$

where $R_m$ is the mitotic radius. See Ref. (16) for additional details.

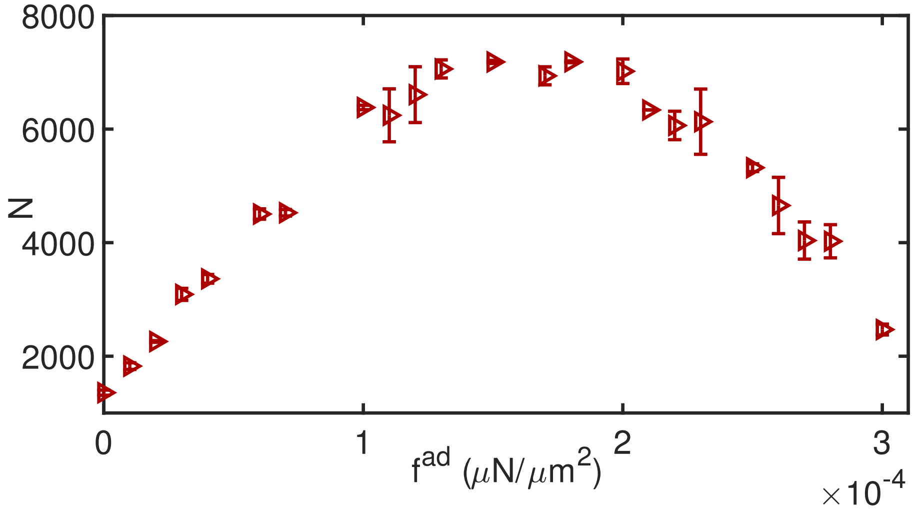


FIG. S1: Number of cells after 7.5 days of growth as a function of cell-cell adhesion strength at $p_c = 5 \times 10^{-5}$ MPa, which is a factor of two less than the value (see Table S1) used in the Main Text.

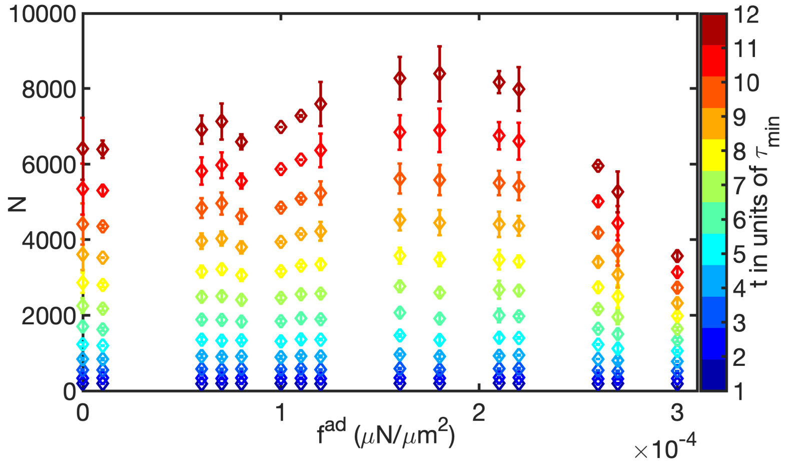


FIG. S2: Number of cells, at $t = \tau_{min}$ to $9\tau_{min}$ as a function of $f^{ad}$. We used $p_c = 1.5 \times 10^{-7}$MPa in the simulations with the Irving-Kirkwood definition of pressure Eq. S.1. These simulations also show that the observed non-monotonic dependence of proliferation on $f^{ad}$ is robust.

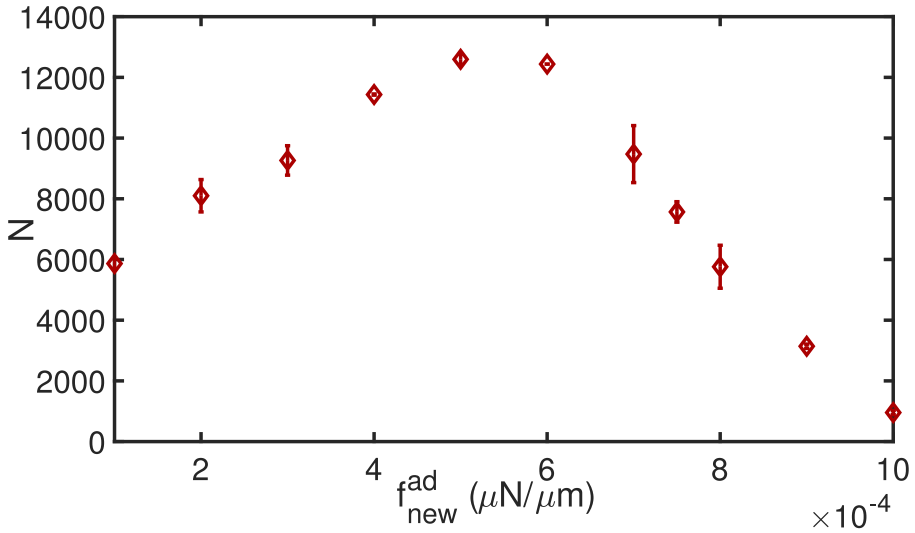


FIG. S3: Number of cells after 7.5 days of growth as a function of cell-cell adhesion strength using an alternate form of $F^{ad} = \rho_m W_s h_{ij}$. The results are qualitatively similar to the ones in Fig. 2 of the Main Text.

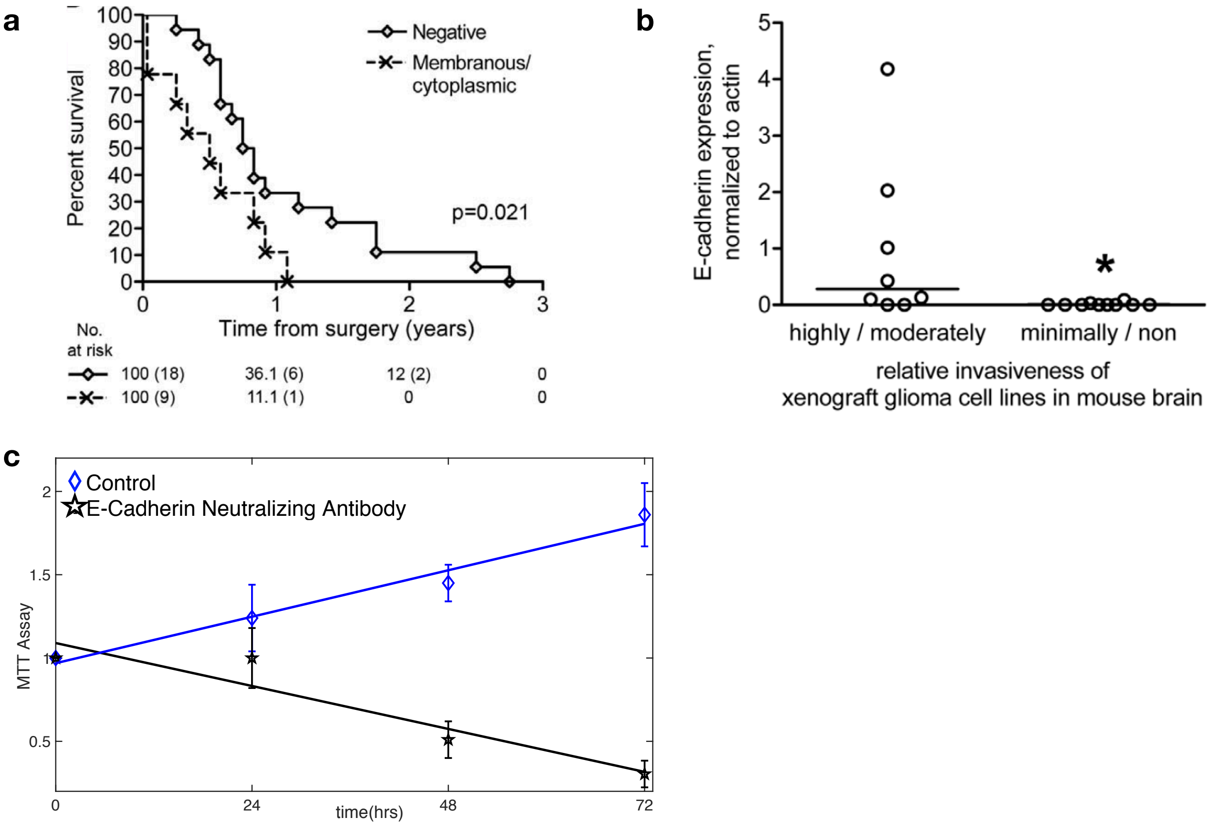


FIG. S4: Figs. **a)** & **b)** are reproduced from Ref. (8). **a)** Overall survival from surgery for patients with tumors characterized by absence (Negative), presence (Membranous/cytoplasmic) E-cadherin expression. Patients whose tumors did not express E-cadherin had better overall survival compared to those that did express E-cadherin. Percent survival (No. of patients) are indicated at the bottom at different times from surgery. **b)** Quantification of E-cadherin expression level compared to Actin versus relative invasiveness of xenograft GMB tumor in mouse brain. **c)** MTT Assay data for Ovarian tumor cells showing proliferation behavior for control cells expressing E-cadherin versus E-cadherin neutralized cells. The experimental results reproduced here give qualitative support to the simulation results.

| **Parameters** | **Values** | **References** |
|---|---|---|
| Critical Radius for Division ($R_m$) | 5 $\mu$m | (17) |
| Extracellular Matrix (ECM) Viscosity ($\eta$) | 0.005 kg/($\mu$m s) | (18) |
| Benchmark Cell Cycle Time ($\tau_{min}$) | 54000 s | (19–21) |
| Adhesive Strength ($f^{ad}$) | $0 - 3 \times 10^{-4} \mu$N/$\mu$m$^2$ | (17), This paper |
| Mean Cell Elastic Modulus ($E_i$) | $10^{-3}$MPa | (18) |
| Mean Cell Poisson Ratio ($\nu_i$) | 0.5 | (17) |
| Death Rate ($b$) | $10^{-6}$s$^{-1}$ | This paper |
| Mean Receptor Concentration ($c^{rec}$) | 0.90 (Normalized) | (17) |
| Mean Ligand Concentration ($c^{lig}$) | 0.90 (Normalized) | (17) |
| Adhesive Friction $\gamma^{max}$ | $10^{-4}$kg/($\mu$m$^2$ s) | This paper |
| Threshold Pressue ($p_c$) | $10^{-4}$MPa | (17, 22) |

TABLE I: The values of the parameters used in the simulations.

**Movies:** In order to visualize the dynamic behavior of pressure experienced by cells, we generated movies from the simulations. All the movies show the three dimensional growth of cell collectives over $\approx$ 7.5 days. Indicated in the color scale with low pressures (blue) and high pressures (red) is the pressure experienced by a cell in units of MPa. They demonstrate vividly the intercellular pressure fluctuations as the cells collectively expand. Each movie frame is spaced at 1000 seconds. The movies have been sped up by a factor of $2.16 \times 10^4$, to aid visualization. The cell cycle time $\tau = \tau_{min}$. Pressure relaxation of cells can be observed as the flickering of colors, indicating cells relaxing from high to low pressure or vice versa. Simulation movies can be found here https://utexas.box.com/s/sl43kcoptciht61y4fm6e4klkagvtv5f

**Movie S1:** Intercellular pressure behavior at $\mathbf{f^{ad} = 0}$: Colormap indicates the intercellular pressure. Cell division and death events are explicitly depicted. Supplementary Movie S1A shows the cross section through the cell collective at $f^{ad} = 0$. Spatial pressure distribution shows elevated pressure in the interior which decreases towards the periphery.

**Movie S2:** Pressure fluctuations in a growing cell collective at intermediate cell-cell adhesion strength $\mathbf{f^{ad} = 1.5 \times 10^{-4}}\mu\text{N}/\mu\text{m}^2$: Merging of two cell spheroids into a larger one can be observed. Growth of single cells and division events are depicted. Low pressure neighborhoods (depicted by blue color) is distributed throughout the surface of the spheroid. Supplementary Movie S2A shows the cross section view at intermediate $f^{ad}$.

**Movie S3:** Intercellular pressure behavior at high cell-cell adhesion, $\mathbf{f^{ad} = 3 \times 10^{-4}}\mu\text{N}/\mu\text{m}^2$: Pressure relaxation behavior for a growing cell collective at high cell-cell adhesion is visualized. Birth, apoptosis, growth and movement of cells is readily observed. Due to high cell-cell adhesion, groups of cells tend to form tightly packed clusters. Merging of such clusters can be seen. Supplementary Movie S3A shows the cross section view at high $f^{ad}$. Pressure experienced by the cells decay as the tumor periphery is approached.

---

[1] JH Irving and John G Kirkwood. The statistical mechanical theory of transport processes. IV. The equations of hydrodynamics. *The Journal of chemical physics*, 18(6):817–829, 1950.

[2] Jerry Zhijian Yang, Xiaojie Wu, and Xiantao Li. A generalized Irving–Kirkwood formula for the calculation of stress in molecular dynamics models. *The Journal of chemical physics*, 137(13):134104, 2012.

[3] Ignacio Ramis-Conde, Dirk Drasdo, Alexander RA Anderson, and Mark AJ Chaplain. Modeling the influence of the E-cadherin-$\beta$-catenin pathway in cancer cell invasion: a multiscale approach. *Biophysical journal*, 95(1):155–165, 2008.

[4] Geert Berx and Frans Van Roy. The E-cadherin/catenin complex: an important gatekeeper in breast cancer tumorigenesis and malignant progression. *Breast Cancer Research*, 3(5):289, 2001.

[5] Fausto J Rodriguez, Laura J Lewis-Tuffin, and Panos Z Anastasiadis. E-cadherin's dark side: possible role in tumor progression. *Biochimica et Biophysica Acta (BBA)-Reviews on Cancer*, 1826(1):23–31, 2012.

[6] DS Tan, HW Potts, AC Leong, CE Gillett, D Skilton, W Hetal Harris, RD Liebmann, and AM Hanby. The biological and prognostic significance of cell polarity and E-cadherin in grade I infiltrating ductal carcinoma of the breast. *The Journal of pathology*, 189(1):20–27, 1999.

[7] Celina G Kleer, Kenneth L van Golen, Thomas Braun, and Sofia D Merajver. Persistent E-cadherin expression in inflammatory breast cancer. *Modern Pathology*, 14(5):458, 2001.

[8] Laura J Lewis-Tuffin, Fausto Rodriguez, Caterina Giannini, Bernd Scheithauer, Brian M Necela, Jann N Sarkaria, and Panos Z Anastasiadis. Misregulated E-cadherin expression associated with an aggressive brain tumor phenotype. *PloS one*, 5(10):e13665, 2010.

[9] Satoshi Utsuki, Yuichi Sato, Hidehiro Oka, Benio Tsuchiya, Sachio Suzuki, and Kiyotaka Fujii. Relationship between the expression of E-, N-cadherins and beta-catenin and tumor grade in astrocytomas. *Journal of neuro-oncology*, 57(3):187–192, 2002.

[10] Carla Perego, Cristina Vanoni, Silvia Massari, Andrea Raimondi, Sandra Pola, Maria Grazia Cattaneo, Maura Francolini, Lucia Maria Vicentini, and Grazia Pietrini. Invasive behaviour of glioblastoma cell lines is associated with altered organisation of the cadherin-catenin adhesion system. *Journal of Cell Science*, 115(16):3331–3340, 2002.

[11] Junji Gofuku, Hitoshi Shiozaki, Toshimasa Tsujinaka, Masatoshi Inoue, Shigeyuki Tamura, Yuichiro Doki, Shigeo Matsui, Shoichiro Tsukita, Nobuteru Kikkawa, and Morito Monden. Expression of E-cadherin and $\alpha$-catenin in patients with colorectal carcinoma: correlation with cancer invasion and metastasis. *American journal of clinical pathology*, 111(1):29–37, 1999.

[12] S Dorudi, JP Sheffield, R Poulsom, JM Northover, and IR Hart. E-cadherin expression in colorectal cancer. An immunocytochemical and in situ hybridization study. *The American journal of pathology*, 142(4):981, 1993.

[13] Karin Sundfeldt, Yael Piontkewitz, Karin Ivarsson, Ola Nilsson, Pär Hellberg, Mats Brännströ, Per-Olof Janson, Sven Enerbäck, Lars Hedin, et al. E-cadherin expression in human epithelial ovarian cancer and normal ovary. *International journal of cancer*, 74(3):275–280, 1997.

[14] Pradeep Reddy, Lian Liu, Chong Ren, Peter Lindgren, Karin Boman, Yan Shen, Eva Lundin, Ulrika Ottander, Miia Rytinki, and Kui Liu. Formation of E-cadherin-mediated cell-cell adhesion activates AKT and mitogen activated protein kinase via phosphatidylinositol 3 kinase and ligand-independent activation of epidermal growth factor receptor in ovarian cancer cells. *Molecular Endocrinology*, 19(10):2564–2578, 2005.

[15] Sun A Kim, Kentaro Inamura, Mai Yamauchi, Reiko Nishihara, Kosuke Mima, Yasutaka Sukawa, Tingting Li, Mika Yasunari, Teppei Morikawa, Kathryn C Fitzgerald, et al. Loss of CDH1 (E-cadherin) expression is associated with infiltrative tumour growth and lymph node metastasis. *British journal of cancer*, 114(2):199, 2016.

[16] Abdul N Malmi-Kakkada, Xin Li, Himadri S Samanta, Sumit Sinha, and Dave Thirumalai. Cell growth rate dictates the onset of glass to fluidlike transition and long time superdiffusion in an evolving cell colony. *Physical Review X*, 8(2):021025, 2018.

[17] Gernot Schaller and Michael Meyer-Hermann. Multicellular tumor spheroid in an off-lattice Voronoi-Delaunay cell model. *Physical Review E*, 71(5):051910, 2005.

[18] Jörg Galle, Markus Loeffler, and Dirk Drasdo. Modeling the effect of deregulated proliferation and apoptosis on the growth dynamics of epithelial cell populations in vitro. *Biophysical journal*, 88(1):62–75, 2005.

[19] James P Freyer and Robert M Sutherland. Regulation of growth saturation and development of necrosis in EMT6/Ro multicellular spheroids by the glucose and oxygen supply. *Cancer research*, 46(7):3504–3512, 1986.

[20] Joseph J Casciari, Stratis V Sotirchos, and Robert M Sutherland. Variations in tumor cell growth rates and metabolism with oxygen concentration, glucose concentration, and extracellular pH. *Journal of cellular physiology*, 151(2):386–394, 1992.

[21] Jacques Landry, James P Freyer, and Robert M Sutherland. Shedding of mitotic cells from the surface of multicell spheroids during growth. *Journal of cellular physiology*, 106(1):23–32, 1981.

[22] Fabien Montel, Morgan Delarue, Jens Elgeti, Laurent Malaquin, Markus Basan, Thomas Risler, Bernard Cabane, Danijela Vignjevic, Jacques Prost, Giovanni Cappello, et al. Stress clamp experi-

ments on multicellular tumor spheroids. *Physical review letters*, 107(18):188102, 2011.
